# Experiment-free learning of exoskeleton assistance remains an unsolved problem

**DOI:** 10.64898/2026.04.01.715109

**Authors:** Steven H. Collins, Friedl De Groote, Robert D. Gregg, He (Helen) Huang, Tommaso Lenzi, Massimo Sartori, Gregory S. Sawicki, Jennie Si, Patrick Slade, Aaron J. Young

**Author notes:** **Correspondence and requests for materials** should be addressed to Steven H. Collins,. Authors listed in alphabetical order by surname; roles described in Author Contributions.

## Abstract

In "Experiment-free exoskeleton assistance via learning in simulation", Luo et al. [1] present an ambitious framework for developing exoskeleton controllers through reinforcement learning exclusively in computer simulation. The authors report that a control policy trained on a small dataset from one subject was directly transferred to physical hardware, reducing human metabolic cost during walking, running, and stair climbing by more than any prior device. If confirmed, this would represent a major breakthrough for the field of wearable robotics and their clinical applications. However, a close examination of the published materials casts doubt on these claims. The reported experimental results violate physiological limits on the relationship between mechanical power and muscle energy use during gait^2,3,4^. The algorithmic claims are surprising and cannot be verified; in contrast with established replicability standards in machine learning^5,6^, executable code has not been made available. We conclude that the goals of this study have not yet been verifiably achieved and make recommendations for avoiding publication errors of this type in the future.

## Main Text

The potential benefits of exoskeleton assistance are limited by human physiology. During gait, joints in the legs cyclically produce and absorb mechanical work, which requires muscles to consume metabolic energy. To produce 1 Joule of positive mechanical work, skeletal muscle consumes about 4 Joules of metabolic energy^2^. Tendons can also store and return energy, which reduces the energy required from muscles. As a result, during gait, 1 Joule of positive joint work is associated with between 2 and 4 Joules of metabolic energy consumption^3,7,8^. This well-known relationship has been incorporated into models that predict the maximum metabolic energy savings achievable with an exoskeleton based on its mechanical power output^4,9^. In practice, imperfections in device control policies and energy transfer to the body reduce these savings. Prior exoskeletons, including those experimentally optimized for individual users, have not exceeded 2.3 Joules of metabolic energy savings per 1 Joule of device work (Figure 1; Supplementary Material).

**Fig. 1:**
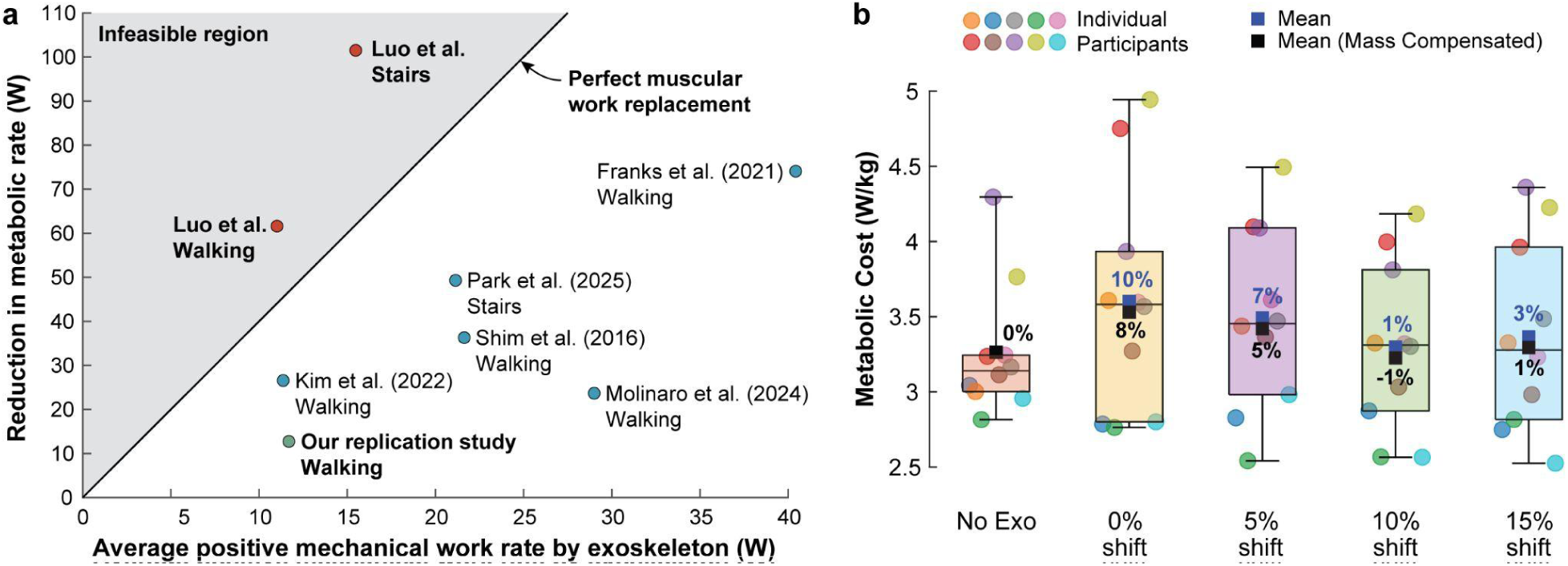
Plausibility of the reported metabolic cost reductions. **a.** Across-study comparison of metabolic energy savings versus positive mechanical work. The y-axis depicts energy savings with respect to wearing the device in zero-torque mode, *i.e.*, the benefits of exoskeleton assistance. The maximum benefit is achieved when all positive exoskeleton work replaces positive muscle work. Two of three conditions from Luo et. al. exceed this theoretical upper limit. Our replication study of the walking condition resulted in a physiologically plausible energy ratio and much lower metabolic energy savings. For reference, we also include best-in-class hip exoskeletons with a variety of control architectures, all of which report energy ratios far from the infeasible region. **b.** Results from our replication study. Participants (N=10) were of similar height, body mass, and age to those in the Luo et al. study and walked at the same speed. We tested the same torque profile on a hip exoskeleton across four different timing conditions to account for potential differences in heel contact detection between studies. No conditions resulted in a statistically significant reduction in metabolic energy cost compared to the no-exoskeleton condition, even when accounting for the slightly greater mass of the exoskeleton used in our tests.

The work and energy values reported by Luo et al. violate these physiological limits. The reported exoskeleton torque profile implies that 1 Joule of positive exoskeleton work resulted in metabolic energy savings of 5.5 Joules during walking and 6.6 Joules during stair climbing (Supplementary Material). These values far exceed both theoretical limits and prior exoskeleton results; we cannot conceive of a physiologically plausible explanation for these data.

We attempted to reproduce the claimed metabolic cost reduction of 24.3% during walking using a similar hip exoskeleton. The feedback control policy from Luo et al. was not reported, so we applied the average resulting torque trajectory, which we expected to produce similar results during steady-state treadmill walking. Our study (Supplementary Materials) was conducted at Harvard University under the same walking conditions (1.25 m/s). Ten healthy participants (N=10) completed five randomized walking conditions: a no-exoskeleton condition and four assistance conditions in which the exoskeleton torque profile was shifted by 0%, 5%, 10%, and 15% of the gait cycle to account for possible differences in heel strike detection timing. Our custom hip exoskeleton had a similar construction and mass (4.8 kg) as the device used by Luo et al. Metabolic rate data were collected using field-standard equipment and methods.

Contrary to Luo et al.’s reported results, this replication found no statistically significant reduction in metabolic cost. The best-performing profile (10% gait cycle shift) resulted in an estimated 1% reduction compared to no exoskeleton (after accounting for the lower mass of the Luo et al. device). No participant had energy savings approaching 24.3% in any condition. Participant metabolic energy cost was reduced by 1.1 Joules for each 1 Joule of positive device work, consistent with theoretical limits and prior hip exoskeletons^10^. We conclude that this is a more reasonable estimate of the metabolic effects of the Luo et al. assistance.

The learning-in-simulation method that forms the centerpiece of the Luo et al. study cannot be replicated from the information provided. The musculoskeletal model, simulation code, reinforcement-learning implementation, and trained policies have not been made available. Only simplified, non-executable pseudocode for reinforcement learning is provided. Important implementation parameters, such as reward function and random seed, are missing (Supplementary Material). The reported results lack necessary evidence of convergence^11^. The authors have consistently declined requests to share the code used to generate their results^12^. This violates reproducibility standards for machine learning in the life sciences^6^, which require the provision of ’the actual computer code’ ^5^ in order for results to be ’worthy of trust’ ^6^.

The algorithmic claims made by Luo et al. are surprising and cannot be verified. Transferring learned policies to real systems is notoriously difficult^13^, requiring large amounts of high-quality data, excellent system models, and significant computation^14^. Luo et al. claim to have created a transferable, task-agnostic policy based on eight gait cycles from one participant in each of three conditions, which did not include task transitions or exoskeleton use; a simplified model of the human that did not include a metabolic model or a nervous system; and a relatively small and rapidly trained neural network. By contrast, recent attempts to learn only musculoskeletal control, without an assistive device, for simpler models of the human body, have required orders of magnitude more trainable nodes, training time, and training data, including task-specific data from many different subjects^15^ (Supplementary Material).

Analysis of the supplementary movies from Luo et al. suggests that human musculoskeletal control was not well captured in the trained model. The described learning-in-simulation approach attempts to learn human musculoskeletal control in parallel with exoskeleton control, thereby simultaneously identifying complementary policies. Musculoskeletal control was not evaluated directly, but renderings of the model that appear in the supplementary movies show infeasible ground reaction forces, discontinuities in segment kinematics, including some that seem to indicate a reset of the simulation, and irregular patterns of torque during transitions between tasks (Supplementary Materials). These issues further undermine confidence in the reported learning-in-simulation results.

Taken together, the implausible claims and lack of transparency of the Luo et al. study, along with our contradictory replication results, strongly suggest that the authors’ goals were not realized. We recommend that researchers with similar interests address deficiencies in both the learning methods (training data, model features, and statistical performance) and model evaluation (kinematics, kinetics, and muscle activity) and carefully apply established experimental methods for measuring physiological outcomes. While we expect that successful wearable robot research will rely heavily on human experiments in the near term, we share the hope that simulation-based approaches will one day streamline the development process.

This example demonstrates the need for more rigorous practices in the publication of human-robot interaction studies. Authors should diligently and dispassionately evaluate each claim, using established methods, and provide sufficient information for complete replication. Editors should include independent reviewers who are experts in the application domain and should require the above standards be met at the time of publication. Together we can maintain scientific integrity as the first consideration in the publication process, earning trust in our works and institutions.

## Supporting information

Mechanical Power Calculation Data and Code

Replication Study Data and Code

## Data availability

All study data necessary to replicate this work are available in the Source Data included with this paper.

## Code availability

Code that reproduces the analysis and results found in Figure 1 and the Supplementary Material is provided as Supplementary Data 1.

## Author Contributions

All authors contributed to the conceptualization, drafting, and revising of the manuscript and gave final approval. S.C. led wordsmithing. F.D.G. led the analysis of replicability of the musculoskeletal model. R.G. led outreach to the original authors. H.H. led the analysis of biomechanical outcomes and contributed to the analysis of the replicability of the learning-in-simulation approach. T.L. contributed to the analysis of biomechanical outcomes. M.S. led the analysis of computational efficiency. G.S. contributed to the analysis of apparent efficiency. J.S. led the analysis of replicability of the learning-in-simulation approach. P.S. conducted the replication study, led the analysis of the replication study, and created code for reproducing this analysis. A.Y. led the analysis of energy ratios, led the across-study comparison, and created code for reproducing these analyses.

## Additional information

### Supplementary information

The online version contains supplementary material available at [].

## Supplementary Material

Accompanies: Experiment-free learning of exoskeleton assistance remains an unsolved problem

### 1. Analysis of Positive Exoskeleton Work and Changes in Metabolic Energy Cost

#### Biological Efficiency of Human Movement

Human locomotion is an efficient process that leverages muscle work, elastic energy storage in tendons, and passive dynamics. In steady, level gait, the body acts like an inverted pendulum; during each step, kinetic and potential energy exchange helps save energy^16^. Even so, muscles and tendons must perform some positive mechanical work to perform tasks such as redirecting the center of mass during step-to-step transitions^17^. Isolated muscle preparations generate positive mechanical work with an efficiency of 0.25; that is, 1 Joule of positive mechanical muscle work consumes 4 Joules of metabolic energy^2,18^. Joint work differs from muscle work, because energy storage and return from spring-like tendons that are in series with muscles also contribute to joint work. During gait, the contribution of tendons can be substantial, thereby reducing the amount of joint work attributed to muscles and increasing the effective ’efficiency’ of joint work. The ’apparent efficiency’ of biological joint work during human walking, defined as the mechanical work produced at joints divided by the net metabolic cost, has been well studied and has values between 25% and 50% ^3,7^. Prior research indicates that the hip joint tends to operate at the lower end of this range, due to its reliance on muscles to perform the vast majority of the mechanical work, unlike the ankle, which captures and returns energy through the Achilles tendon structure and tends to have higher apparent efficiency^17^.

#### Apparent Efficiency and Energy Ratio in Exoskeleton Assistance

The effectiveness of exoskeleton work in reducing metabolic energy use can be quantified in a similar manner. This has often been referred to as ‘apparent exoskeleton efficiency’ or ’transfer efficiency’ in the literature^17,4,19^. Numerous studies have used this measure, which is formally the positive mechanical work performed by the exoskeleton on the human divided by the net human metabolic cost reduction due to the use of the exoskeleton. Here, the change in metabolic energy cost is calculated relative to the cost of walking with the exoskeleton in a zero-torque mode, so as to isolate the effects of assistance from the costs related to the added mass of the exoskeleton. Typically, these analyses are only performed on the positive mechanical work produced by the exoskeleton, for two reasons. First, most exoskeleton controllers are tuned to maximize positive mechanical work while providing very little negative work. That is, they try to avoid resisting the user, because this is typically neither preferred nor beneficial. Second, the cost to a muscle of performing concentric (positive) work is much higher than that of performing eccentric (negative) work in terms of muscle metabolism^20^. Apparent exoskeleton efficiency can also be reported as an equivalent ’energy ratio’ of mechanical work or power to the corresponding change in metabolic energy or power, depicted as ’mechanical:metabolic’. For example, an apparent efficiency of 0.25 corresponds to an energy ratio of 1:4.

#### Achievable Values of Exoskeleton Energy Ratio

Energy ratio is a useful measure for understanding how much an exoskeleton reduces metabolic cost with a given assistance profile from an assistive device acting in parallel to a joint. A larger energy ratio indicates a larger metabolic cost reduction per unit mechanical input from the exoskeleton. The most advantageous energy ratio possible would be a value of 1:4, which would mean that for each 1 Joule of positive mechanical work provided by the exoskeleton, 1 Joule of positive muscle work is replaced and a reduction of 4 J in metabolic energy expenditure is achieved for the human. Based on the known efficiencies of walking and human muscle, this is the best energy ratio achievable in theory. However, this has not been achieved previously for a number of reasons. First, while baseline apparent efficiencies of human joints can be as low as 0.25, this value represents the lower limit, and empirically determined values of apparent efficiency for joints in the lower limbs during locomotion are always higher, often much higher^3^. When an exoskeleton assists a joint with a higher apparent efficiency, the potential for positive exoskeleton work to reduce human metabolic cost is lower. Second, exoskeleton control policies are always imperfect, and sometimes completely ineffective, such that positive work done by the exoskeleton does not maximally reduce human muscular effort, and can even cause it to increase. Third, energy transfer through an exoskeleton interface with the human body is imperfect. There are always losses due to, for example, compliance and damping in the soft tissues of the human body. This final loss is consistent with the second law of thermodynamics, which indicates that energy transfer will always induce some level of entropy in the form of heat, which results in energy loss. Thus, there is good reason to believe that an ideal energy ratio of 1:4, positive mechanical work to metabolic energy savings, is not achievable in practice.

#### Estimation of the Positive Mechanical Power of the Luo et al. Exoskeleton during Walking

We downloaded the supplemental data provided by Luo et al. to obtain the N = 8 subject-specific device hip torques collected during level walking at 1.25 m/s. These data appear in cf. Fig. 3 of the original manuscript and are depicted in fig. S1 below. Because Luo et al. did not include kinematics data from the human or exoskeleton, we instead downloaded an open-source dataset^21^ which contained N=22 able-bodied subjects with age and body mass that closely matched those of the participants in the Luo et al. study. For this analysis, we assumed that hip exoskeleton assistance had little effect on hip joint kinematics, which is supported by prior studies in which able-bodied individuals walked with hip exoskeletons at similar speeds and magnitudes of assistance^22,23,24^. We extracted the hip joint kinematics including hip angles and velocities from these data at the most similar level walking speed (1.2 m/s) and calculated a subject-averaged hip joint velocity normalized to the gait cycle (fig. S1). Each subject’s individual torque profile was multiplied by this hip angular velocity curve to generate a subject-specific exoskeleton joint power curve (fig. S1). Negative portions of the curve were set to zero so as to isolate positive mechanical power. The average value of the curve was then calculated. This value was multiplied by 2, because the system was bilateral, yielding an average positive exoskeleton joint power for each subject. Across all participants, the average positive mechanical power from the Luo et al. exoskeleton during walking was estimated to be 11.2 ± 2.0 W (mean ± st. dev.).

**Supplementary figure S1:**
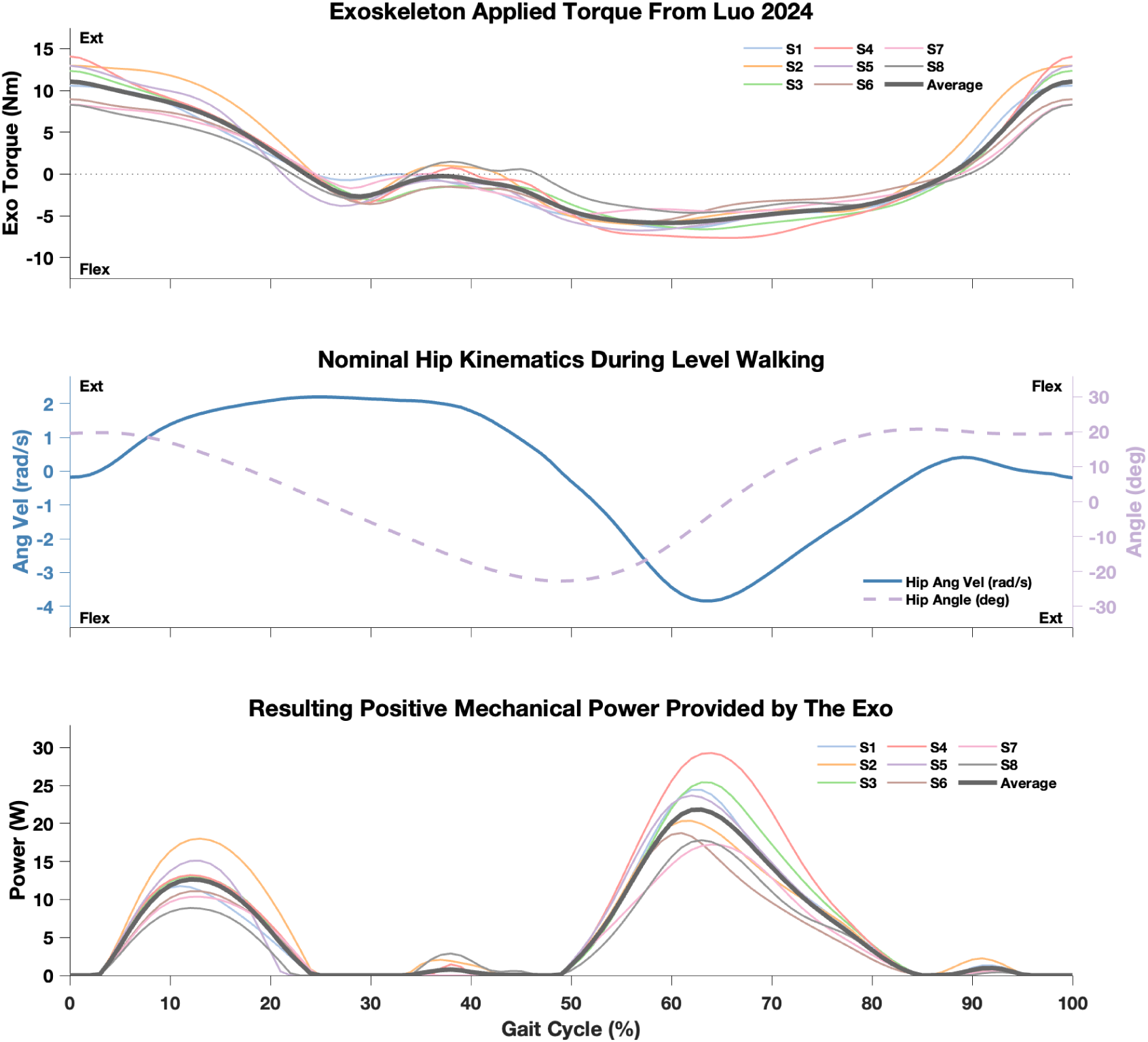
Visualization of the exoskeleton mechanical power calculation. *Top:* Data from Luo et al. showing the applied exoskeleton torque on a per subject basis, as well as the averaged torque over a gait cycle. *Middle:* Data from an open-source dataset^21^ showing human hip joint angles and velocities during walking at 1.2 m/s. *Bottom:* Calculated mechanical power provided by the exoskeleton, obtained by multiplying the applied torques by the hip angular velocities. Only positive mechanical power is included.

#### Calculating the Energy Ratio of the Luo et al. Exoskeleton during Walking

We divided the average reduction in metabolic energy cost of 61.6 W reported by Luo et al. for the walking condition by our estimate of the average positive mechanical power of the Luo et al. exoskeleton in the same condition. The resulting energy ratio was 1:5.5, or 1 W positive mechanical exoskeleton power yielding 5.5 W metabolic energy savings for the human. This value is substantially greater than the theoretical limit of 1:4. We cannot conceive of a physiologically plausible reason for this discrepancy.

#### Analyzing the Sensitivity to Exoskeleton Power Assumptions

To account for potential differences in heel strike timing between the Luo et al. study and the study from which hip joint velocity were taken, we shifted the kinematic data by plus and minus 5% of the gait cycle in 1% increments, across a 10% window, and calculated the resulting positive mechanical work at each increment. We expected this range to encompass all reasonable possibilities for synchronizing the kinetic and kinematic data. This resulted in a range of mechanical powers from 9.4 W to 13.8 W and a range of energy ratios from 1:4.5 to 1:6.5. That is, across all reasonable possibilities of torque and velocity timing, the torque and metabolic rate data reported by Luo et al. imply a physiologically inexplicable benefit to the user.

#### Verification Based on Replication Study Data

Our calculations of the mechanical power of the Luo et al. device are limited by the fact that kinematics data were not reported. While changes in hip joint velocity due to exoskeleton assistance are typically small, they could affect the positive work done by the exoskeleton. Fortunately our replication study, described in detail below, provided us with internally consistent kinetics and kinematics data for calculation of power and validation of the above approach. We applied the average torque profile from the Luo et al. study using a similar hip exoskeleton during treadmill walking at 1.25 m/s among N = 10 subjects with similar characteristics. We calculated the average positive mechanical power using data from exoskeleton torque and velocity sensors, and found that the applied mechanical power of the hip exoskeleton ranged between 11.1 W and 13.0 W, with an average value of 11.7 W. This closely matches the outcome of our calculation using the reported exoskeleton torque and open-source kinematics, which yielded an average value of 11.2 W. Thus, two separate means of estimating the positive mechanical power of the exoskeleton in the Luo et al. study produce very similar results. This increases our confidence in the calculated energy ratios presented here.

#### Calculation of Energy Ratios for the Running and Stair Climbing Conditions

We also calculated the exoskeleton energy ratios for running and stair climbing, this time using data from cf. Figure 4 in the Luo et al. study, downloaded as supplementary materials. In this N = 1 analysis (only a single representative subject was provided in the article for these conditions), we took the steady-state portion of each plot, integrated the positive portions of the power curve, and divided by the corresponding time. There were a total of 4 strides that could be selected during running at 2.0 m/s and 6 strides during stair climbing. Within each condition, all strides had similar average positive power. The average rate at which mechanical work was produced by the exoskeleton was found to be 63.5 W during running and 15.5 W during stair climbing. Using the metabolic rate data reported by Luo et al. for the respective activities, this resulted in an energy ratio of 1:1.9 (mechanical:metabolic) during running and 1:6.6 during stair climbing. As with the level walking condition, the energy ratio during stair climbing is well outside the plausible range and corresponds to nearly three times as much metabolic energy reduction per unit mechanical device work as the best prior result (Figure 1a).

#### Comparison of Energy Ratios Across Exoskeleton Walking Studies

We quantified the energy ratios for a number of state-of-the-art exoskeleton studies that report large reductions in metabolic rate during walking, some of which were carried out by authors of this perspective (Figure 1a; table S1). We focus on hip exoskeleton studies for this analysis to enable the most direct comparison to the Luo et al. study. Studies that performed tailored tuning and optimization on a per-subject basis using human-in-the-loop optimization (HILO) provided substantially greater metabolic benefits per unit mechanical power compared to generalized controllers that did not change control parameters across individuals. We analyzed Seo et al. (2016), which led to the development of the Samsung GEMS device. This system produced a large reduction in metabolic cost and obtained an energy ratio of 1:1.7, mechanical:metabolic, the best-in-the-field for non-HILO results. We also analyzed Molinaro et al. (2024), which used a data-driven neural network similar to that used by Luo et al., and obtained an energy ratio of 1:0.8. Our replication study of Luo et al. obtained a similar energy ratio of 1:1.1. A comparison of the torque patterns in these three studies can be found in figure S2. Franks et al (2021) performed HILO at the hip during walking, applying very large torques using a tethered emulator system, and obtained an energy ratio of 1:1.8. With much smaller torques and an autonomous, soft exosuit optimized to individual users, Kim et al. (2022) obtained an energy ratio of 1:2.3. This is the best such ratio for hip assistance during level walking, and it is only slightly better than half of the theoretical maximum energy ratio of 1:4. In contrast, the data provided in Luo et al. indicates an energy ratio of 1:5.50. This is ∼37% above the theoretical maximum and more than double the energy ratio of the best-in-class results previously published in the field. Luo et al. did not use a personalized model or tuning through human-in-the-loop optimization, which was necessary to achieve best-in-class results previously. Notably, Luo et al. do not claim that their method should outperform HILO in terms of benefits to the user, but instead only in terms of the development and experimental time required to obtain effective assistance patterns. They provide no rationale as to why their approach should yield a ratio of device mechanical work to metabolic energy savings that exceeds physiological limits and the results of prior devices. Details for the calculation of each of these outcomes are provided in table S1.

#### Comparison of Energy Ratios Across Exoskeleton Stair Climbing Studies

Unlike level-walking, the metabolic benefits of using hip exoskeletons during stair climbing have not been demonstrated frequently in the literature. This may be because stair ascent with an exoskeleton has rarely been tested, or because it is difficult to obtain metabolic measurements under these conditions^27^. One of the few examples is reported by Park et al. (2025), which performed HILO of hip assistance and found an energy ratio of 1:2.33, mechanical to metabolic. This assistance only reduced metabolic cost by 4% relative to the no exoskeleton condition, because the mass penalty of wearing the exoskeleton during stair ascent is especially large. The Luo et al. exoskeleton caused a similar metabolic penalty during the zero torque condition, provided lower values of torque assistance, and did not tune assistance to individual participants, yet is claimed to have led to larger metabolic energy savings and three times the energy ratio.

**Supplementary table S1.** Data used in our analysis of the apparent efficiency of different hip exoskeletons. We selected studies that reported large reductions in metabolic cost, which would be expected to have the lowest apparent efficiencies and largest energy ratios. All studies used hip exoskeletons on able-bodied adults. Five studies addressed walking, two addressed stair ascent, and one addressed running. Three studies included human-in-the-loop optimization of exoskeleton assistance, in which exoskeleton control parameters were tuned so as to maximize the reductions in metabolic rate on an individual basis. For each study, we report: the average, positive mechanical power from the exoskeleton; the corresponding reduction in metabolic rate for the human compared to wearing the exoskeleton while it applied zero-torque; the resulting ’apparent efficiency’ of the exoskeleton, which we define as the change in metabolic rate divided by the positive power supplied by the exoskeleton; and the ’energy ratio’, which we define as the ratio of average positive mechanical power from the exoskeleton to the reduction in metabolic rate (mechanical:metabolic). Well-established physiological limitations set the theoretical lower bound on apparent efficiency at 0.25 and the corresponding upper bound on energy ratio at 1:4. Only the Walking and Stair ascent conditions reported by Luo et al. violate these limits; no values from any other study analyzed approach these limits, including our replication study.

| Study | Joint | Task | HILO | Exoskeleton power (W) | Metabolic change (W) | Apparent efficiency | Energy ratio |
| --- | --- | --- | --- | --- | --- | --- | --- |
| Seo et al. (2016) | Hip | Walking | No | 21.6 | 36.3 | 0.60 | 1:1.7 |
| Franks et al. (2021) | Hip | Walking | Yes | 40.4 | 74.1 | 0.55 | 1:1.8 |
| Kim et al. (2022) | Hip | Walking | Yes | 11.4 | 26.6 | 0.43 | 1:2.3 |
| Molinaro et al. (2024) | Hip | Walking | No | 29.0 | 23.7 | 1.22 | 1:0.8 |
| Park et al. (2025) | Hip | Stairs | Yes | 21.1 | 49.3 | 0.43 | 1:2.3 |
| Luo et al. (2024) | Hip | Walking | No | 11.2 | 61.6 | 0.18 | 1:5.5 |
|  | Hip | Running | No | 63.5 | 112.7 | 0.56 | 1:1.8 |
|  | Hip | Stairs | No | 15.5 | 101.5 | 0.15 | 1:6.6 |
| Our replication study | Hip | Walking | No | 11.7 | 12.7 | 0.92 | 1:1.1 |

**Supplementary figure S2.**
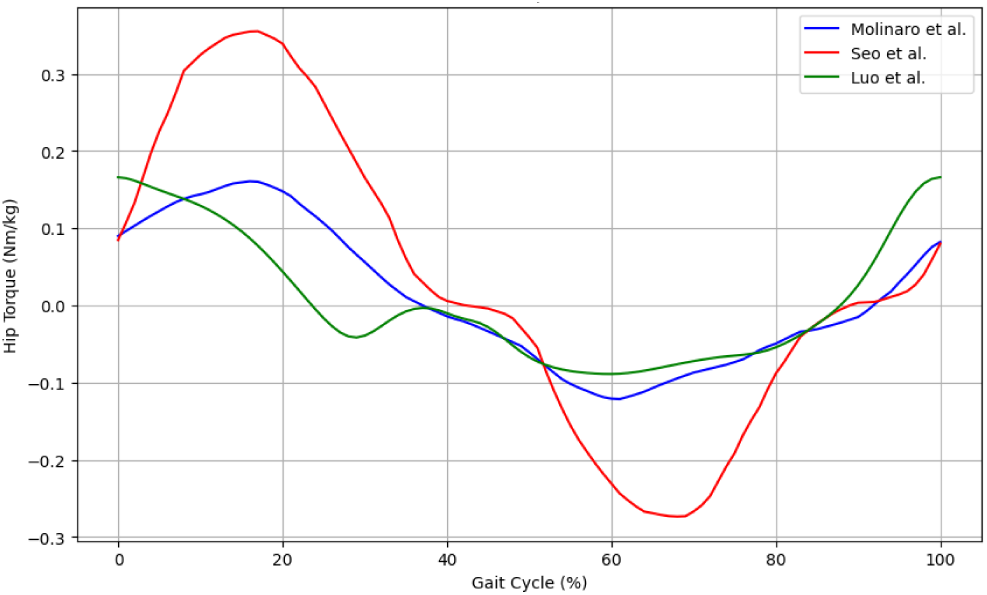
Examples of torque patterns from prior studies of hip exoskeleton assistance during level walking. The pattern of torque from Luo et al. (green) is similar to that from Molinaro et al. (blue), yet Luo et al. reports about three times greater reduction in metabolic rate (62 W vs. 24 W). Seo et al. (red) applied a torque with about twice as much peak torque as in Luo et al., yet also found a reduction in metabolic rate that was about half the value from Luo et al. (36 W vs. 62 W). These results are consistent with the energy ratio analysis.

#### Limitations in the Energy Ratio Analysis

One potential limitation of this analysis is that only positive mechanical work was considered. Negative mechanical work can also contribute to reductions in metabolic cost. For the studies considered, however, the negative work done by the active hip exoskeletons was negligible; the Luo et al. study reported that negative work comprised only 3.5% of the total mechanical work. For passive exoskeletons^29,30^, the negative work is typically larger than the positive work (as much as 2 times the positive mechanical work), and cannot be less than the positive mechanical work. For these systems, and for activities in which human negative work dominates (such as steep declines) the negative work contributions can become non-negligible. Note that, in the above analyses, we consider positive and negative work separately, and not net work over a cycle; even passive devices typically provide substantial positive work by this approach. Additionally, in static tasks where muscles must hold a force isometrically, such as holding a squat, there is a metabolic cost associated with isometric force production that cannot be explained based on mechanical joint work (which could be zero). During gait, however, the cost of force production is strongly tied to mechanical work production in the joints because the joints are simultaneously moving. Importantly, in cases where muscle force production is thought to provide a better explanation of metabolic rate than work production, the expected benefits of 1 Joule of exoskeleton assistance are <u>lower</u> than if muscle work costs dominated. For example, during slow walking, in which the calf muscles hold the Achilles tendon nearly isometrically and consume metabolic energy primarily to produce an isometric force, the benefits of one unit of ankle exoskeleton work, or torque, are expected to be lower than during fast walking or stair ascent, in which the calf muscles produce more positive work.

### 2. Methods of the Replication Study

#### Study Protocol

Participants (n = 10, age = 23 ± 3 yrs, height = 1.69 ± 0.07 m, weight = 70 ± 16 kg, mean ± st. dev.) completed a series of conditions to evaluate the torque assistance profile during level walking on a treadmill at 1.25 m/s. Most participants were novice exoskeleton users. Participant characteristics reported by Luo et al. for their walking study were similar (n = 8, age = 27 ± 3 yrs, height = 1.72 ± 0.09 m, weight = 68 ± 19 kg). The experience level of the participants with the hip exoskeleton was not stated in Luo et al., however there was no training protocol described in the paper, which suggests the users may have been novices. The assistance conditions in our replication study included normal walking (no exo) and walking with torque assistance. During assisted conditions, the exoskeleton provided the average torque profile for level walking at 1.25 m/s reported by Luo et al. Four assistance conditions were evaluated with the torque profile shifted to start at slightly different parts of the gait cycle, to account for potential differences in heel strike detection between our device and the device used in the original study. These four conditions included starting the torque profile at the instant of estimated heel strike (0% change in gait cycle phase) and three conditions in which the profile was delayed by some percentage of the gait cycle duration (5%, 10%, and 15%). Participants completed 5 minutes of walking on the treadmill to warm up followed by 5 minutes of quiet standing and then the experimental conditions. The ordering of the 5 experimental conditions were randomized and presented in a double-reversal order (ABCDEEDCBA) to account for any ordering effects. Each condition was 5 minutes in length. Metabolics were recorded with a COSMED K-5 portable respirometry system, which measured both oxygen consumption and carbon dioxide production. The steady-state metabolic cost was estimated by taking the average over the last 2 minutes of respirometry data from each condition, averaging across presentations, and applying a field-standard equation^31^. This calculation differs from the approach of Luo et al., in which carbon dioxide production was not measured.

#### Potential Differences in Heel Strike Estimation

The heel strike detection in the replication study relied on a phase variable estimated from a thigh-worn IMU that evaluates the thigh angle and sagittal plane angular velocity^32^. The heel strike for this IMU-based estimation occurs when the thigh sagittal plane angular velocity changes direction, which for level ground walking occurs slightly before the heel hits the ground. Luo et al. estimated a phase variable from the learned models. Since these models are not available and ground-truth data from ground reaction forces are not available with the torque profiles, the timing of their phase estimation and subsequent detection of heelstrikes is unclear. We experimentally evaluated four assistance conditions using the same torque profile shifted by a percentage of the gait cycle duration to account for potential differences in timing.

#### Pilot Testing that Informed the Experimental Protocol

A pilot study with a single participant initially evaluated three different torque profiles across the distribution of the participants reported in Luo et al.: the mean across all participants, the mean plus one standard deviation, and the mean minus one standard deviation. Each of these torque profiles were similar (within a few Nm of torque) since the profiles across participants had little variation. These torque profiles resulted in similar metabolic costs (within 2% of each other). Due to the similarity of these profiles and metabolic results, we selected to only evaluate the mean torque profile for the full experiment with ten participants. Another pilot experiment with a single participant evaluated many possible gait cycle shifts to apply to the mean torque profile. These conditions included -10% to 30% shifts of the heelstrike timing at increments of 5% of the gait cycle. The four conditions from 0% to 15% of the gait cycle were selected for the full experiment because they had the lowest corresponding metabolic energy cost.

#### Hip Exoskeleton Details

This experiment was performed with a custom hip exoskeleton (fig. S3). The exoskeleton began as an M-BLUE hip exoskeleton design^33^ and was modified to be able to provide higher torques. The hip exoskeleton weighs 4.8 kg, which includes the weight of the batteries necessary for the hour-long walking experiments. This system has the same architecture as the device used by Luo et al. and is very similar in its details.

**Supplementary figure S3.**
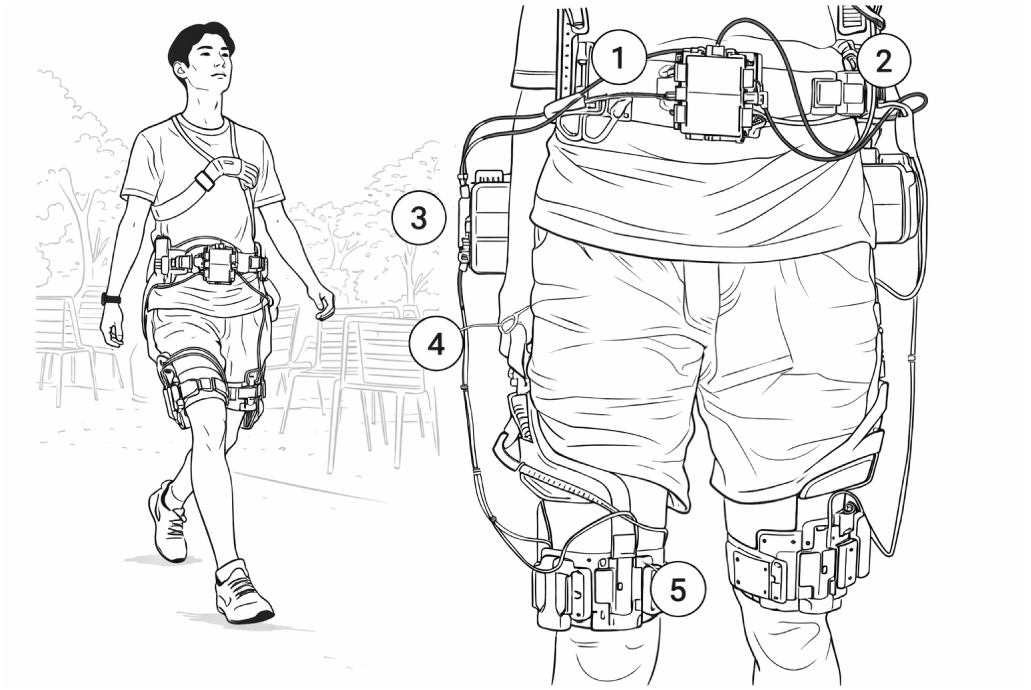
The custom hip exoskeleton used to perform the replication experiment. The exoskeleton assists hip flexion and extension by applying hip torque using (1) a microcontroller and (2) portable batteries to drive (3) brushless motors (AK10-9; CubeMars). Torque is measured by (4) load cells and transmitted to the body by (5) custom carbon fiber braces on the pelvis and thigh. While the exoskeleton is depicted outside in the photograph, all tests were conducted during steady, treadmill walking. (Illustration to meet bioRxiv requirements.)

#### Metabolic Cost Adjustment for Different Hip Exoskeleton Weights

The hip exoskeleton in the replication study weighed 4.8 kg, while the exoskeleton tested by Luo et al. had a reported mass of 3.2 kg. We adjusted the metabolic cost in the replication experiments to account for the 1.6 kg lower mass of the exoskeleton used in the original experiment, providing a reduction in the estimated metabolic cost from the replication study. We estimated the reduction in metabolic cost that corresponds to reducing the mass of the hip exoskeleton by 1.6 kg using the relationship identified in a prior experimental study^34^. That study found that 1 kg of mass added to the hips increased metabolic rate by 0.045 W/kg. The 1.6 kg difference in exoskeleton mass would therefore be expected to lead to a difference of 0.072 W/kg in metabolic rate. We subtracted this value from the metabolic rate we measured to provide an improved estimate of the energy cost of walking with the Luo et al. device.

#### Experimental Results

On average, participants increased metabolic cost during all the evaluated assistance conditions compared to not wearing the exoskeleton (Figure 1b). The best performing timing of the assistance profile was when the torque profile was delayed by 10% of the gait cycle. Adjusting for the difference in the weight of their exoskeleton and our device, the best condition would result in a 1% reduction in metabolic cost compared to not wearing the exoskeleton. There were no statistically significant reductions between any assistance condition and the no exoskeleton condition. The single best metabolic reduction for any participant during any exoskeleton condition was approximately 14%. These results strongly differ from the average metabolic reduction of 24.3% reported by Luo et al.

#### Results with Non-Responders Withheld

Two participants (red and olive circles in Figure 1b) had an increase of greater than 1 W/kg in each assistance condition compared to not wearing the exoskeleton. While it is not uncommon for some participants to experience an increase in metabolic rate when generic exoskeleton torques are applied, especially if those torques are not particularly beneficial, it is possible that other factors could have contributed to the increase in this case, such as inadequate training. In order to provide the most generous analysis possible, we recalculated our study average without these two participants. Omitting these data, the 10% gait cycle shifted torque provided estimated metabolic energy savings of 6% compared to not wearing the exoskeleton, after accounting for the lower mass of the Luo et al. device. That is, even with a bias due to post-hoc participant selection, the result still strongly differs from that reported by Luo et al. We do not endorse the practice of removal of participant data from study-level analysis after finding that their results were less favorable than other participants.

#### Mechanical Power

Exoskeleton data were recorded to compute mechanical power applied by the exoskeleton. These data include the applied exoskeleton torque measured by a force sensor, the thigh angle estimated from the motor encoder position, and the thigh-worn IMU data for each condition.The average positive mechanical power was computed from the applied exoskeleton torque and sagittal-plane thigh angular velocity data. We then calculated the average positive mechanical power for each assistance condition (fig. S4). All conditions resulted in similar mechanical power. The 0% and 5% shifted assistance conditions resulted in the highest power, but did not correspond to the largest reductions in metabolic cost. That is, the apparent efficiency in these conditions was higher and the energy ratio lower than in the 10% shift condition. The raw and processed exoskeleton data used to compute the average positive mechanical power and other power-related metrics are available as Supplementary Data.

**Supplementary figure S4.**
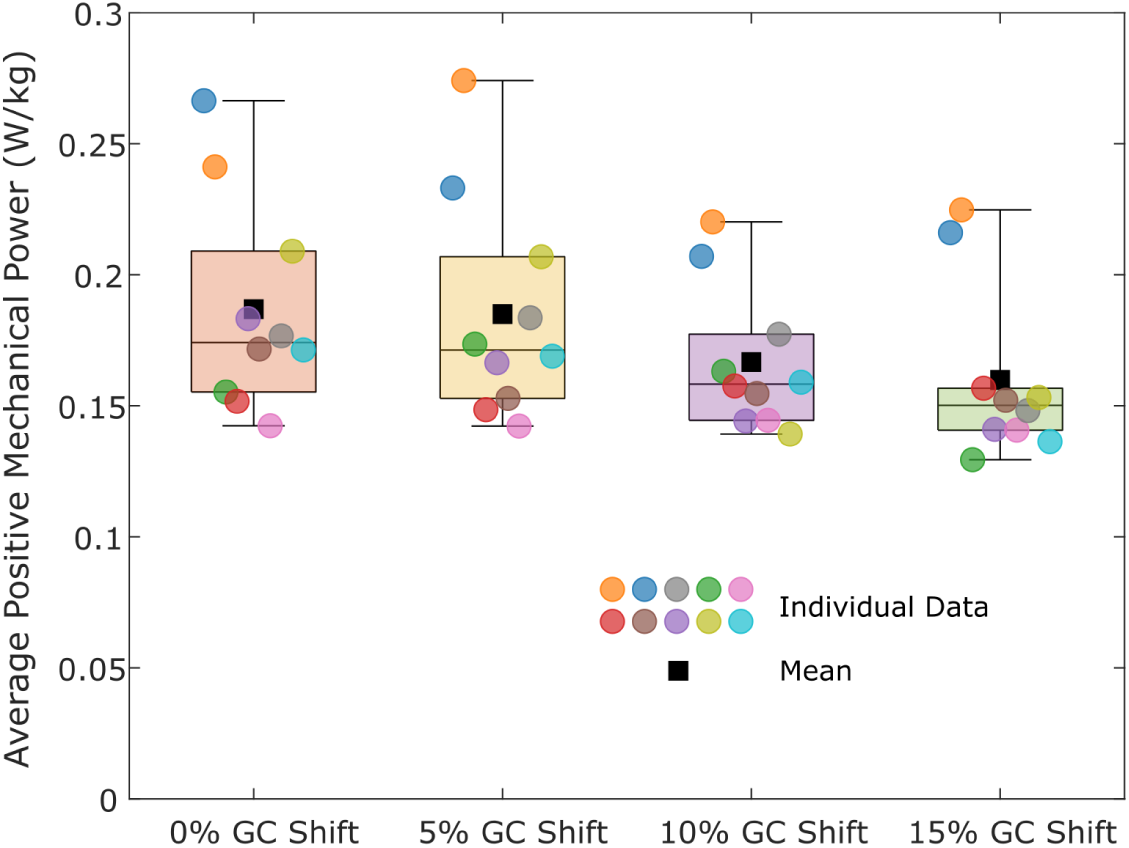
The average positive mechanical power of the different experimental conditions tested in our replication study during 1.25 m/s treadmill walking with the average torque profile from Luo et al. shifted by 0%, 5%, 10%, or 15% of the gait cycle (GC).

#### Limitations of our Replication Study

While we made every effort to replicate the walking experiment described by Luo et al., some aspects of the comparison are imperfect. We sought to test the exact control policy used by Luo et al., which set desired torque as a function of the recent history of measured thigh angles. Unfortunately, the trained policy was not published and not made available upon request. We were therefore only able to test the average resulting torque trajectory as a function of time normalized to percent stride. In general, we would expect well-designed state feedback control policies to be more responsive to the needs of the user on a stride-to-stride basis, potentially providing more effective assistance. However, during the steady treadmill walking conditions tested here, we expect such differences to be muted. It should also be noted that differences in control architecture would not explain the differences in energy ratio that we observed, because torques arising from any policy manifest in the measured torque and power used to calculate the energy ratio. Prior studies using similar forms of learned feedback control, such as Molinaro et al. (2024), produced energy ratios similar to that in our replication study and strongly different from the Luo et al. study (table S1). Another limitation is that we did not test a zero-torque condition, but instead estimated the energy cost of such a condition based on models of the effects of added mass. Fortunately, these models have been found to accurately predict the cost of added mass in many exoskeleton studies^9^. Another limitation is that there could be differences between the timing of the application of torque between the two studies. We addressed this factor by testing several candidate timing values and reporting results for the most effective condition. Prior studies of the effects of subtle changes in the timing of hip exoskeleton assistance suggest that small differences in timing could not have had a substantial impact on our results^23,35^. While all of these imperfections are worth consideration, none could plausibly account for the large differences between the results reported by Luo et al. and those found in our replication study. We observed a reduction in metabolic rate that was nearly five times smaller than the value reported by Luo et al. (13 W vs. 62 W), corresponding to a percent change of 1% vs. 24% compared to normal walking. Prior replications of the benefits of exoskeleton assistance have revealed much smaller differences. For example, Slade et al. (2022) found energy savings of 32% for an ankle exoskeleton control policy that resulted in energy savings of 31% in Poggensee et al. (2021). As another example, Nuckols et al. (2020) found energy savings of 4% using a tethered ankle exoskeleton emulator to deliver rotational stiffness similar to an unpowered elastic exoskeleton that resulted in savings of 7% in Collins et al. (2015).

### 3. Analysis of the Replicability of the Learning-in-Simulation Approach

A learning-in-simulation method forms the primary intellectual contribution of the Luo et al. study. Unfortunately, Luo et al. omit several parameters that are necessary to define the reinforcement learning problem for the human model and the exoskeleton controller, making the related claims unreproducible. The learning signal is not fully specified, leaving the controller without a clear objective or a basis for performance optimization; important parameters related to the reward function are not disclosed and the reward computation is absent from the pseudocode. The exoskeleton controller training procedure based on Proximal Policy Optimization (PPO) is also methodologically incomplete; no training curves, loss trajectories, convergence data, or evaluation steps are presented, and the pseudocode omits important PPO settings necessary for implementation. Consequently, the reported convergence and performance outcomes cannot be independently verified and remain empirically unsubstantiated.

#### Undisclosed Parameters in the Reward Formulation

Reward formulation is crucial in reinforcement learning. Reward design information is needed for reproducibility, comparison to other methods, and interpretability. Specifically, to reproduce published results, readers need the exact reward function, including all terms, coefficients, scaling/normalization, and any shaping. This information is typically provided in the form of a downloadable and executable code repository that produces the results reported in the publication. Unfortunately, no code is provided by Luo et al. Instead, an incomplete description of the approach is provided in the main text, supplementary materials, and pseudocode. In the Methods section, Luo et al. define the following reward function that forms the exoskeleton control objective for reinforcement learning.

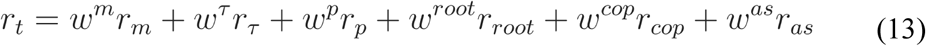

While the five weights and three reward terms are defined in Equation 13, the two sub-rewards (joint-position error *r_p_* and pelvis-position error *r_root_* ), are not sufficiently defined because the parameters *σ_p_* for sub-reward *r_p_* (Equation 7) and *σ_root_* for sub-reward *r_root_* (Equation 8) are not disclosed. Consequently, it remains unclear how the two positional terms are incorporated, and to what degree, in the reinforcement learning objective to optimize control performance.

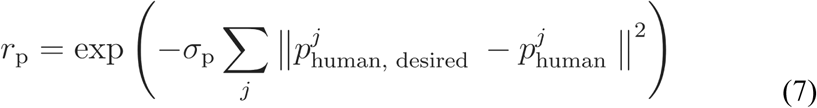

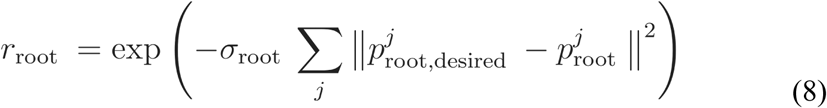

Consequently, it remains unclear what roles these two positional terms play in the formulated reinforcement learning objective used to optimize control performance. Omitting the scale parameters *σ_p_* and *σ_root_* effectively leaves the reward function undefined; these two terms are important for shaping gait kinematics, and without them there is significant uncertainty when attempting to reproduce the intended control behavior. If the values of these weights are low, the learned policy may deviate substantially from the target movement patterns during optimization. This could lead to reward hacking^39^, which occurs when the reward or objective function is misspecified, incomplete, or misaligned with the true intent, allowing the agent to exploit loopholes in the reward signal and discover spurious strategies that do not genuinely solve the intended task. For example, in a recent study applying reinforcement learning to a soft exosuit^40^, a systematic ablation analysis demonstrated that removing the kinematic error term from the learning objective caused the controller to fail to maintain normative walking patterns, leading to elevated EMG activity and less effective assistance. Without a complete code stack that includes these and other terms, it is not possible to replicate the learning process and verify that it produces the described results.

#### Incomplete Exoskeleton Controller Training Procedure based on PPO

Luo et al. use Proximal Policy Optimization (PPO) to train the exoskeleton controller. Unfortunately, the description of the training procedure is incomplete and therefore unreproducible. Several hyperparameters in the PPO implementation are not provided, including those that define rollout length, gradient clipping, random seeds, and normalization of the observation and reward. Actor structure details, such as the activation functions and the output distribution and parametrization, are also not provided. The referenced reward functions in the pseudocode (GetExoReward and GetHumanReward) are absent from the provided environment file, meaning the learning signal is missing and the PPO controller cannot be trained or evaluated from the supplied materials. No analysis of run-to-run variability in learning is provided, and there is no indication as to whether multiple seeds were used. No evaluation during training was provided, such as deterministic evaluation rollout, evaluation logging with learning performance metrics, or held-out test episodes to evaluate the final policy. No learning curves, convergence data, or evaluation steps were provided; all reported performance results, *e.g.*, torque tracking and metabolic-rate reductions, stem from post-training deployment. The stopping criteria were defined informally as “when the reward value of the exoskeleton control network … stopped increasing (around iteration 3,500)” without a precise definition of patience or tolerance. For these reasons, the PPO reinforcement learning methodology lacks the transparency and methodological rigor necessary to support the major algorithmic claims or to ensure scientific reproducibility^5,6^.

### 4. Reproducibility of the Musculoskeletal Model

An important component of the learning-in-simulation method is the musculoskeletal model. Unfortunately, Luo et al. do not provide the model nor the information that would be needed to build the model, compromising the reproducibility of the study. The standard in musculoskeletal modeling is to share models, e.g. through platforms such as SimTK or with the simulation code.

#### Musculoskeletal Model

It is unclear whether a previously published model was used. Luo et al. only provide a high level description without specifying parameters describing inertial body segment properties, joint axis/center locations, and muscle properties. The authors mention that the model has 8 revolute joints and 14 ball-and-socket joints but list body segments (except for shoulder) rather than joints. Reference cf. 53 in the Methods section of the Luo et al. paper describes a similar model but although this model has the same number of joints, it has a different number of muscles. There are no other references to papers describing similar models in the section describing the model. The authors provide an expression for the passive muscle force length curve in cf. Eq. 3 (without specifying the parameters), but not for the active muscle force length curve or force velocity curve. It is not clear whether these were obtained from Thelen et al. (ref. 54 in the main manuscript). It is unclear how tendons were modelled, for example whether they were modeled as rigid or elastic and, if elastic, what tendon stiffnesses were used. There is no description of the lever arms or wrapping geometries for muscle-tendon units. There is no description of how the interaction between the feet and the ground was modelled. Parameters that define body geometry, segment mass properties, muscle mechanics, tendon mechanics, and contact mechanics all have a strong effect on the behavior of musculoskeletal simulations. For example, tendon and foot-ground contact stiffness can have a large influence on simulated joint kinematics and powers^41^. Many different choices are possible, illustrated by a wide variety of model layouts and parameter values in the literature. Without these specifications, it is not possible to create a reasonable approximation of the Luo et al. musculoskeletal model.

#### Model of Human-Exoskeleton Interface

The model of the human-exoskeleton interface is described in some detail and parameters are given in Supplementary Table 1. Yet, crucial information is lacking. It is unclear which reference frames were used to compute translations between body segments and exoskeleton (d in eq. 5 of the Luo et al. paper) and the parameters provided do not have units.

#### Ambiguous Definitions

Confidence in rigor of the modeling approach is undermined by inaccuracies in describing the model. Some symbols in cf. Supplementary Table 8 do not match symbols in the Methods section. The authors write that ‘muscle activations … determine the change of the muscle fiber length and further, the change of muscle force’, which is imprecise because any given activation can result in different shortening velocities or levels of force depending on the operating conditions of the muscle. In some locations f_M_ is defined as tension, whereas in others it seems to be a normalized force (unitless), for example in cf. Eq. 2, in which f_M_ is multiplied by the maximum isometric force to get muscle force.The Methods section states that the motion imitation network produces joint angle profiles but this does not agree with cf. Fig. 2c, in which the output of the network seems to be joint torques. If executable code had been provided, minor communication errors such as these would be of less importance, but without code they do impede replicability.

### 5. Modeling and Computational Efficiency Comparison

Luo et al. make strong claims regarding the computational efficiency of their learning-in-simulation approach. These claims cannot be verified because executable code has not been provided. We sought to test the reasonableness of these claims by comparing them to a recent study, Simos et al. (2025), that sought to fulfill only the human motor control imitation portion of the objectives of the Luo et al. study. We chose this comparison study because it also uses imitation learning and Proximal Policy Optimization, and because it verifies not only kinematic similarity across a variety of human movements, but also tests that the joint kinetics and muscle activity underlying those movements match human values from the training set. The comparison study used a simpler musculoskeletal model, yet required: a neural network that was orders of magnitude larger; orders of magnitude more training data; and an order of magnitude more computational time. We conclude that the unverified Luo et al. claims would also be surprising compared to benchmark efforts with verifiable results.

#### Comparison of Musculoskeletal Models

Luo et al. report using a full-body musculoskeletal model with 208 muscle-tendon units, 50 degrees of freedom in human joints, and 2 degrees of freedom in exoskeleton joints. They report using HyFyDy as the physics engine. Simos et al. used a comparatively simpler musculoskeletal model. The trunk and lower-limb musculoskeletal model included 80 muscle-tendon units, 20 degrees of freedom in human joints, and no exoskeleton. This is about 2.6 times fewer body states than Luo et al. They used MyoSuite^43^ running on MuJoCo as the physics engine.

#### Comparison of Neural Network Models

Luo et al. describe a control architecture comprising three neural networks representing motion imitation for the human body, muscle imitation for the human, and exoskeleton control. The motion imitation network had 100 inputs, two layers with 256 nodes each, for both actor and critic, and 50 output states for the actor and 1 output state for the critic. The muscle imitation network had 50 input states, three layers with 512, 256, and 256 nodes, and 208 output states. The exoskeleton control network had either 18 input states (extended fig. 4) or 16 input states (Methods subsection on "data-driven learning of exoskeleton control"), two layers with 128 and 64 nodes for both actor and critic, and 2 outputs for the actor and 1 output for the critic. Taken together, the total number of weights and biases for these neural networks is 494,662, assuming fully connected layers with biases. Simos et al. describe a control architecture comprising four neural networks in a mixture-of-experts (MoE) paradigm, with three expert networks and one gate network. Each expert network had 309 input states, six layers with 2,048, 1,536, 1,024, 1,024, 512, and 512 nodes, and 80 output states. The gate network had 309 input states, three layers with 1,024, 512, and 256 nodes, and 3 output states. This results in 22,676,723 weights and biases, or about 46 times the network parameters in Luo et al. That is, the comparison model required two orders of magnitude more network parameters per model degree of freedom to achieve reasonable performance.

#### Comparison of Training Data

The Luo et al. study reports that training data comprised three, 10-second trials of gait data from a single participant, one trial each for steady-state walking, running, and stair ascent conditions. In the comparison study, Simos et al. report requiring 1,084 trials, averaging about 6 seconds in length, for a total of 6,840 seconds of training data. Trials were selected from a public database^44^to address a rich set of behaviors, including walking, running, backwards walking, arc walking, and turning in place, each at a variety of speeds and including non-steady motions, performed by 21 human participants. That is, the comparison study required substantially more diverse training data and two orders of magnitude more training data to train control networks for a substantially simpler musculoskeletal with no assistive device.

#### Comparison of Training Time

The Luo et al. study reports that training the neural networks required 8 hours of clock time on an NVIDIA RTX3090 GPU. In the comparison study, Simos et al. report that training the neural networks required 240 hours (10 days) on a moderately faster GPU, the NVIDIA A100. That is, the comparison study required 30 times the computation time as the Luo et al. study. This difference likely relates to the much larger neural networks required to produce verifiably similar musculoskeletal control in the comparison study.

### 6. Analysis of the Biological Plausibility of the Simulations shown in the Supporting Movies

Luo et al. present a learning-in-simulation approach in which exoskeleton control is learned alongside a model of human motor control using a musculoskeletal model. For the exoskeleton control to be effective, and to transfer to real human users, it is necessary for both the model of human motor control and the musculoskeletal model to be sufficiently realistic. Unfortunately, Luo et al. do not directly evaluate the simulation results; they do not, for example, compare measured and simulated muscle activations, joint kinetics, ground reaction forces, or joint kinematics. As a result, our understanding of the simulation outcomes is limited to what we can interpret from the Supplementary Movies through a frame-by-frame visual analysis. This analysis is sufficient, however, to indicate that there were important discrepancies.

#### Discontinuities in Kinematics

On several occasions, the musculoskeletal simulation seems to change the pose of its body instantaneously. For example, in Supplementary Movie 1 (“MOESM2”), the model appears to begin to fall between frames 22:26 and 22:28. Then, in the next frame at time 22:29, the model has returned to a more typical walking pattern. Such kinematic discontinuities do not occur in human gait, and we are unable to identify an explanation for them.

#### Erroneous Ground Reaction Forces

Interaction forces between the ground and the body have an important effect on the body’s movement and strong implications for the joint moments and muscle forces that were needed to produce those movements. At several instances in the Supplementary Movies, the ground reaction forces deviate substantially from typical values or produce patterns that are not possible while conforming to physical constraints. For example, the Supplementary Movies sometimes show ground reaction forces with a large horizontal component and negligible vertical component (fig. S5). This is not possible for finite values of coefficient of friction; the foot would slip rather than produce a large horizontal force. The videos often depict ground reaction force vectors that rapidly vary in magnitude and direction, unlike human ground reaction forces that smoothly vary over the course of the stance phase^45,46^. The feet of the simulation model often appear to penetrate the ground. It is not clear whether the joint moments, and corresponding muscle forces, in the musculoskeletal simulation are also deviating substantially from normal and physically possible values, or whether there is an error in the physics of the musculoskeletal model. In either case, these erroneous ground reaction forces indicate a significant mismatch between the real and simulated systems, which would be expected to lead to learned control policies that do not transfer well from simulation to experiments with real users.

#### Irregular Patterns of Torque during Transitions

Luo et al. claim a method for generating exoskeleton control across and between a variety of locomotor behaviors. They place an emphasis in the manuscript on smoothly varying exoskeleton torques as the user transitions between behaviors (cf. Fig. 4). However, Supplementary Movie 2 (“MOESM3”) depicts exoskeleton torques with abrupt jumps and irregular fluctuations during transitions between walking and running. This suggests that the learned exoskeleton control policy may not have generalized to movements outside the training set, which comprised a few seconds of data in three steady-state conditions for one participant. This would compound problems associated with ineffective exoskeleton control policies within the conditions of the training set.

**Supplementary figure S5.**
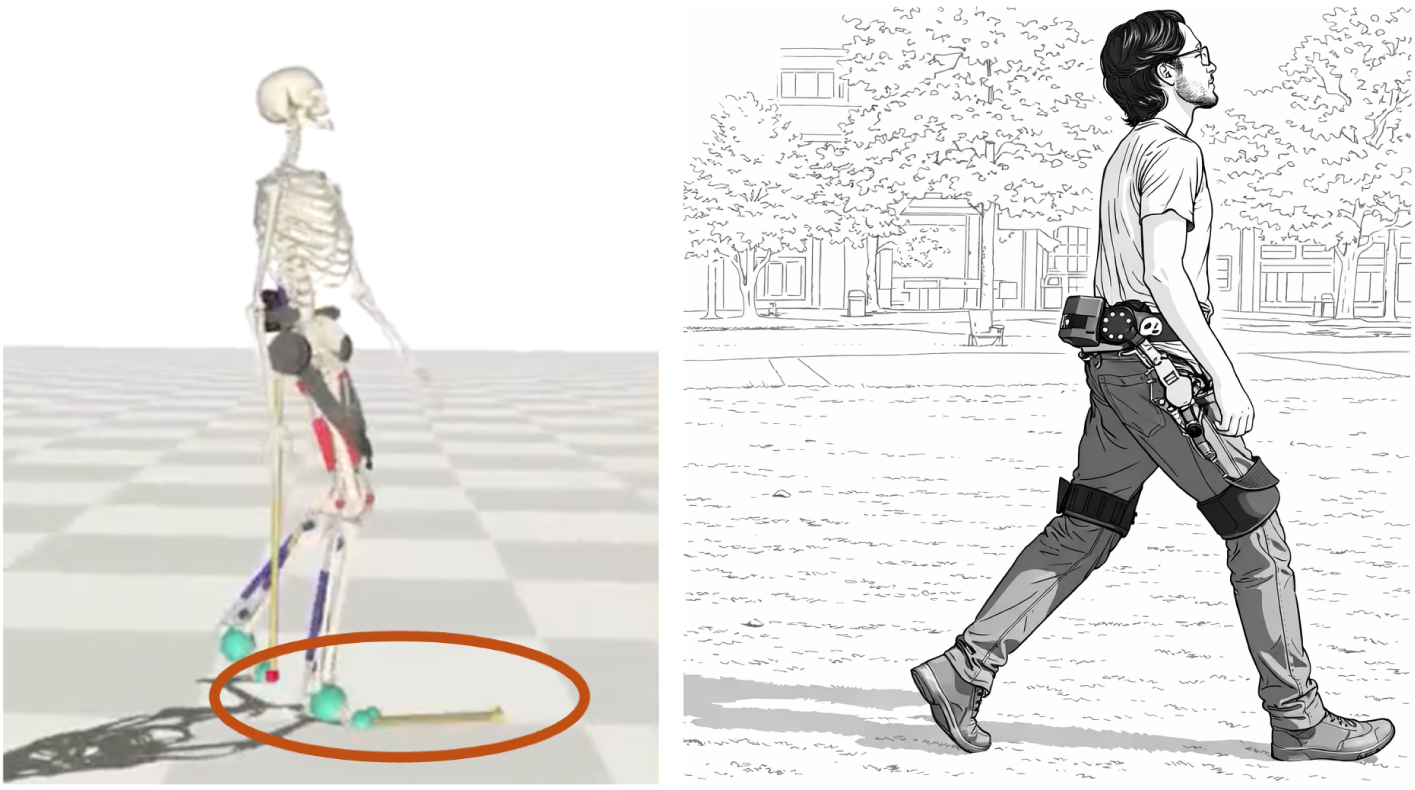
Example of an unrealistic simulation result depicted in the supplementary movies. This frame was captured from Supplementary Movie 2 at timestamp 19:15. In the simulation, the model’s right foot exhibits a physically impossible ground reaction force; for finite coefficient of friction, if the vertical component of force were zero, as shown, the foot would slide rather than generate a large forward ground reaction force. This frame also shows large differences between human body state and that of the musculoskeletal simulation, suggesting the learned human control policy did not capture actual human motor control well. (In this preprint only, to comply with bioRxiv policies, at right we show an illustration of the human body state from the original movie.)

