## Supplementary material for "Experiment-free learning of exoskeleton assistance remains an unsolved problem": Mechanical Power Calculation Data and Code: Plot_Individual_Subjects.m

```
% Required Files:
%   1. 'Luo_Paper_All_Data_update.mat'
%   2. 'hip_flexion_1.2ms-1.mat'
%
%% 0. Initialization and Load Data
clear;
clc;
close all;
disp('Loading data for individual subject analysis (v15)...');

try
    load('Luo_Paper_All_Data_update.mat', 'fig3b_torques_125_all_subjects');
    disp('Loaded ''fig3b_torques_125_all_subjects'' (101x8 matrix).');
catch
    error('Could not find ''Luo_Paper_All_Data_update.mat'' or variable ''fig3b_torques_125_all_subjects''.');
    return;
end

try
    hip_data_file = 'hip_flexion_1.2ms-1.mat';
    hip_data_struct = load(hip_data_file);
    hip_field_names = fieldnames(hip_data_struct);
    hip_angle_deg_1001 = hip_data_struct.(hip_field_names{1});
    disp(['Loaded ''', hip_field_names{1}, ''' (as angle in deg).']);
catch
    error(['Could not find ''', hip_data_file, '''.']);
    return;
end

disp('---');

% --- USER TOGGLE ---
plot_rectified_power = true;
% -------------------

T_gait_cycle = 1.0; % seconds (Still needed for calculation)
n_subjects = 8;
n_points = 101;

% Individual Body Masses (Participants 1-8)
body_masses_kg = [62, 70, 70, 85, 109, 50, 56, 45]; % (1x8 row vector)
disp(['Using individual body masses (n=8): ', num2str(body_masses_kg), ' kg']);

TITLE_FONT_SIZE = 18;
LABEL_FONT_SIZE = 16;
TICK_FONT_SIZE = 14;
LEGEND_FONT_SIZE = 11;
AVG_LINE_WIDTH = 3.5;
INDIV_LINE_WIDTH = 1.5;
AVG_LINE_COLOR = [0.4 0.4 0.4]; % Dark gray for mean line
ANG_VEL_COLOR = [70, 130, 180]/255; % "Steel Blue"
AXIS_LABEL_FONT_SIZE = 12;

% Define 8 pastel colors
pastel_colors = [
    174, 199, 232; % Light Blue
    255, 187, 120; % Light Orange
    152, 223, 138; % Light Green
    255, 152, 150; % Light Red
    197, 176, 213; % Light Purple (*** WILL BE USED FOR ANGLE ***)
    196, 156, 148; % Light Brown
    247, 182, 210; % Light Pink
    150, 150, 150  % Medium Gray
] / 255; % Normalize to 0-1

PASTEL_PURPLE = pastel_colors(5, :);

%% Prepare Angular Velocity (Static, 101x1)
disp('Step 2: Preparing static hip angular velocity profile...');
x_original_phase = linspace(0, 100, length(hip_angle_deg_1001));

% This is now our primary X-axis variable
x_target_phase_percent = linspace(0, 100, n_points)'; % (101x1) column

hip_angle_deg_101 = interp1(x_original_phase, hip_angle_deg_1001, x_target_phase_percent, 'spline');
d_theta_d_phase_deg = gradient(hip_angle_deg_101, 1.0);
dp_dt = 100 / T_gait_cycle;
hip_ang_vel_deg_s = d_theta_d_phase_deg * dp_dt;
hip_ang_vel_rad_s = deg2rad(hip_ang_vel_deg_s);

ang_vel_ext_is_pos_rad_s = -hip_ang_vel_rad_s(:); % (101x1) column
ang_vel_ext_is_pos_deg_s = -hip_ang_vel_deg_s(:); % (101x1) column


hip_angle_flex_is_pos_deg = hip_angle_deg_101(:); % (101x1) column

disp('  Static velocity profile (Pos=Extension) created.');

%% Prepare Torque Data (101x8)
disp('Step 3: Preparing 8-subject torque matrix [Nm]...');

torques_ext_is_pos_Nm_kg = fig3b_torques_125_all_subjects; % (101x8) [Nm/kg]
torques_ext_is_pos_Nm = torques_ext_is_pos_Nm_kg .* body_masses_kg; % (101x8) [Nm]
avg_torque_Nm = mean(torques_ext_is_pos_Nm, 2); % (101x1) [Nm]

disp('  8-subject torque matrix [Nm] (Pos=Extension) created.');

%% Calculate Power (101x8)
disp('Step 4: Calculating 8-subject power matrix [W]...');

instantaneous_power_w = torques_ext_is_pos_Nm .* ang_vel_ext_is_pos_rad_s;
avg_instantaneous_power = mean(instantaneous_power_w, 2);
power_neg_as_zero = instantaneous_power_w;
power_neg_as_zero(power_neg_as_zero < 0) = 0;
avg_rectified_power = mean(power_neg_as_zero, 2);

disp('  All power calculations complete.');

%% Final Report (Console)
mean_positive_power_per_subject = mean(power_neg_as_zero, 1);
overall_mean_positive_power = mean(mean_positive_power_per_subject);
fprintf('\n--- INDIVIDUAL SUBJECT RESULTS (v15 - Pos=Ext) ---\n');

for i = 1:n_subjects
    fprintf('  Subject %d (Mass: %.0f kg) Mean Positive Power = %.3f W\n', i, body_masses_kg(i), mean_positive_power_per_subject(i));
end

fprintf('  -------------------------------------------------\n');
fprintf('  Overall Mean Positive (Extension) Power = %.3f W\n', overall_mean_positive_power);
fprintf('-------------------------------------------------\n');

%% Final 3-Panel Plot
disp('Generating final 3-panel plot (v15 - Gait Cycle)...');
figure('Name', 'Individual Subject Analysis (v15)', 'Position', [100, 100, 1000, 900]);

% --- Panel 1: Torque [Nm] ---
ax_torque = subplot(3, 1, 1);
set(ax_torque, 'ColorOrder', pastel_colors, ...
    'FontSize', TICK_FONT_SIZE, 'FontWeight', 'bold');
hold(ax_torque, 'on');

p_indiv_trq = plot(ax_torque, x_target_phase_percent, torques_ext_is_pos_Nm, 'LineWidth', INDIV_LINE_WIDTH);
p_avg_trq = plot(ax_torque, x_target_phase_percent, avg_torque_Nm, 'Color', AVG_LINE_COLOR, ...
    'LineWidth', AVG_LINE_WIDTH, 'DisplayName', 'Average');

title('Exoskeleton Applied Torque From Luo 2024', 'FontSize', TITLE_FONT_SIZE, 'FontWeight', 'bold');
ylabel(ax_torque, 'Exo Torque (Nm)', 'FontSize', LABEL_FONT_SIZE, 'FontWeight', 'bold');
legend(ax_torque, [p_indiv_trq; p_avg_trq], ...
    {'S1', 'S2', 'S3', 'S4', 'S5', 'S6', 'S7', 'S8', 'Average'}, ...
    'Location', 'northeast', 'FontSize', LEGEND_FONT_SIZE, 'NumColumns', 3, 'Box', 'off');
yline(ax_torque, 0, 'k:', 'HandleVisibility', 'off');
add_axis_padding(ax_torque, 0.1, 'left'); % Pad the left axis

yl_torque = get(ax_torque, 'YLim');
text(ax_torque, 1, yl_torque(2), 'Ext', 'VerticalAlignment', 'top', 'FontSize', AXIS_LABEL_FONT_SIZE, 'FontWeight', 'bold');
text(ax_torque, 1, yl_torque(1), 'Flex', 'VerticalAlignment', 'bottom', 'FontSize', AXIS_LABEL_FONT_SIZE, 'FontWeight', 'bold');
set(ax_torque, 'Box', 'off', 'TickDir', 'out', 'XTickLabel', []); % Clean-up

yyaxis(ax_torque, 'right');
set(ax_torque, 'YTick', [], 'YColor', 'none');
yyaxis(ax_torque, 'left'); % Return focus

% --- Panel 2: Angular Velocity [rad/s] & Angle [deg] ---
ax_ang_vel = subplot(3, 1, 2);
set(ax_ang_vel, 'FontSize', TICK_FONT_SIZE, 'FontWeight', 'bold');
hold(ax_ang_vel, 'on');

% Left Y-Axis (Velocity)
yyaxis(ax_ang_vel, 'left');
p_vel = plot(ax_ang_vel, x_target_phase_percent, ang_vel_ext_is_pos_rad_s, ...
    'Color', ANG_VEL_COLOR, 'LineWidth', 2.5, 'DisplayName', 'Hip Ang Vel (rad/s)');
ylabel(ax_ang_vel, 'Ang Vel (rad/s)', 'FontSize', LABEL_FONT_SIZE, 'FontWeight', 'bold');
set(ax_ang_vel, 'YColor', ANG_VEL_COLOR);

add_axis_padding(ax_ang_vel, 0.1, 'left');
yl_vel = get(ax_ang_vel, 'YLim');
text(ax_ang_vel, 1, yl_vel(2), 'Ext', 'VerticalAlignment', 'top', 'FontSize', AXIS_LABEL_FONT_SIZE, 'FontWeight', 'bold');
text(ax_ang_vel, 1, yl_vel(1), 'Flex', 'VerticalAlignment', 'bottom', 'FontSize', AXIS_LABEL_FONT_SIZE, 'FontWeight', 'bold');

% Right Y-Axis (Angle)
yyaxis(ax_ang_vel, 'right');
p_ang = plot(ax_ang_vel, x_target_phase_percent, hip_angle_flex_is_pos_deg, ...
    '--', 'Color', PASTEL_PURPLE, 'LineWidth', 2.5, 'DisplayName', 'Hip Angle (deg)');
ylabel(ax_ang_vel, 'Angle (deg)', 'FontSize', LABEL_FONT_SIZE, 'FontWeight', 'bold');
set(ax_ang_vel, 'YColor', PASTEL_PURPLE);
add_axis_padding(ax_ang_vel, 0.1, 'right');

% Re-add correct labels for the ANGLE axis (Pos=Flex)
yl_ang = get(ax_ang_vel, 'YLim');
text(ax_ang_vel, 99, yl_ang(2), 'Flex', 'VerticalAlignment', 'top', 'HorizontalAlignment', 'right', 'FontSize', AXIS_LABEL_FONT_SIZE, 'FontWeight', 'bold');
text(ax_ang_vel, 99, yl_ang(1), 'Ext', 'VerticalAlignment', 'bottom', 'HorizontalAlignment', 'right', 'FontSize', AXIS_LABEL_FONT_SIZE, 'FontWeight', 'bold');

title('Nominal Hip Kinematics During Level Walking', 'FontSize', TITLE_FONT_SIZE, 'FontWeight', 'bold');
legend([p_vel, p_ang], 'Location', 'northeast', 'FontSize', LEGEND_FONT_SIZE, 'Box', 'off');
set(ax_ang_vel, 'Box', 'off', 'TickDir', 'out', 'XTickLabel', []); % Clean-up

% --- Panel 3: Power [W] ---
ax_power = subplot(3, 1, 3);
set(ax_power, 'ColorOrder', pastel_colors, ...
    'FontSize', TICK_FONT_SIZE, 'FontWeight', 'bold');
hold(ax_power, 'on');

if plot_rectified_power
    p_indiv_pwr = plot(ax_power, x_target_phase_percent, power_neg_as_zero, 'LineWidth', INDIV_LINE_WIDTH);
    p_avg_pwr = plot(ax_power, x_target_phase_percent, avg_rectified_power, 'Color', AVG_LINE_COLOR, ...
        'LineWidth', AVG_LINE_WIDTH, 'DisplayName', 'Average');
    title('Resulting Positive Mechanical Power Provided by The Exo', 'FontSize', TITLE_FONT_SIZE, 'FontWeight', 'bold');
else
    p_indiv_pwr = plot(ax_power, x_target_phase_percent, instantaneous_power_w, 'LineWidth', INDIV_LINE_WIDTH);
    p_avg_pwr = plot(ax_power, x_time_sec, avg_instantaneous_power, 'Color', AVG_LINE_COLOR, ...
        'LineWidth', AVG_LINE_WIDTH, 'DisplayName', 'Average');
    title('Resulting Mechanical Power Provided by The Exo', 'FontSize', TITLE_FONT_SIZE, 'FontWeight', 'bold');
end


xlabel(ax_power, 'Gait Cycle (%)', 'FontSize', LABEL_FONT_SIZE, 'FontWeight', 'bold');

ylabel(ax_power, 'Power (W)', 'FontSize', LABEL_FONT_SIZE, 'FontWeight', 'bold');
legend(ax_power, [p_indiv_pwr; p_avg_pwr], ...
    {'S1', 'S2', 'S3', 'S4', 'S5', 'S6', 'S7', 'S8', 'Average'}, ...
    'Location', 'northeast', 'FontSize', LEGEND_FONT_SIZE, 'NumColumns', 3, 'Box', 'off');
yline(ax_power, 0, 'k:', 'HandleVisibility', 'off');
add_axis_padding(ax_power, 0.1, 'left');

set(ax_power, 'Box', 'off', 'TickDir', 'out'); % Clean-up

yyaxis(ax_power, 'right');
set(ax_power, 'YTick', [], 'YColor', 'none');
yyaxis(ax_power, 'left');


linkaxes([ax_torque, ax_ang_vel, ax_power], 'x');
xlim(ax_torque, [0, 100]); % Set x-axis from 0 to 100%

disp('Final 3-panel plot (v15) generated.');

%% --- Helper Function ---
function add_axis_padding(ax, padding_fraction, axis_side)
    % Adds a symmetrical padding to the specified Y-axis of a plot
    if nargin < 3
        axis_side = 'left'; % Default to left
    end
    
    yyaxis(ax, axis_side);
    current_ylim = get(ax, 'YLim');
    data_min = current_ylim(1); data_max = current_ylim(2);
    
    if isempty(data_min) || isempty(data_max) || ~all(isfinite(current_ylim)), return; end
    if data_min == data_max, data_min = data_min - 0.5; data_max = data_max + 0.5; end
    
    data_range = data_max - data_min;
    padding = data_range * padding_fraction;
    
    if data_min >= 0 && (data_min - padding < 0)
        new_ylim = [0, data_max + padding];
    else
        new_ylim = [data_min - padding, data_max + padding];
    end
    
    set(ax, 'YLim', new_ylim);
end
```
