## Supplementary material for "Experiment-free learning of exoskeleton assistance remains an unsolved problem": Replication Study Data and Code: generateFileNameList.m

```
function fileNameList = generateFileNameList(subjectNum)

S = sprintf('S%d', subjectNum);

folderPath = fullfile(pwd, S);

files = dir(fullfile(folderPath, ['*' S '*']));
if isempty(files)
    error('No files found containing "%s" in folder: %s', S, folderPath);
end

fileNameList = {
    [S ' Exo C1']
    [S ' Exo C2']
    [S ' Exo C3']
    [S ' Exo C4']
    [S ' Exo C4-2']
    [S ' Exo C3-2']
    [S ' Exo C2-2']
    [S ' Exo C1-2']
    };
end
```
