## Supplementary material for "Experiment-free learning of exoskeleton assistance remains an unsolved problem": Replication Study Data and Code: main_all.m

```
clear;

subjectNumList = [1, 2, 3, 4, 5, 6, 7, 8, 9, 10];

for i = 1:length(subjectNumList)

    subjectNum = subjectNumList(i);
    fileNameList = generateFileNameList(subjectNum);
    parts = split(fileNameList{1});
    pathName = parts{1};

    powerList = zeros(length(fileNameList), 1);

    for j = 1:length(fileNameList)
        fileName = fileNameList{j};
        matFileName = sprintf('%s Power.mat', fileName);
        matFullPath = fullfile(pathName, matFileName);

        load(matFullPath);

        powerList(j) = power_pos_avg_norm;
    end

    half_size = length(fileNameList)/2;
    fileNameList = {fileNameList{1:half_size}};
    powerList(1:half_size) = (powerList(1:half_size) + powerList(end:-1:half_size+1)) / 2;
    powerList(half_size+1:end) = [];

    powerListAll(i, :) = powerList(1:half_size);
end

plot_power_all(powerListAll);
```
