## Supplementary material for "Experiment-free learning of exoskeleton assistance remains an unsolved problem": Replication Study Data and Code: main_average_power_metabolics.m

```
clear;

subjectNumList = [1, 2, 3, 4, 5, 6, 7, 8, 9, 10];

for i = 1:length(subjectNumList)

    subjectNum = subjectNumList(i);
    fileNameList = generateFileNameList(subjectNum);
    parts = split(fileNameList{1});
    pathName = parts{1};

    powerList = zeros(length(fileNameList), 1);

    for j = 1:length(fileNameList)
        fileName = fileNameList{j};
        matFileName = sprintf('%s Power.mat', fileName);
        matFullPath = fullfile(pathName, matFileName);

        load(matFullPath);

        powerList(j) = power_pos_avg_norm;
    end

    half_size = length(fileNameList)/2;
    fileNameList = {fileNameList{1:half_size}};
    powerList(1:half_size) = (powerList(1:half_size) + powerList(end:-1:half_size+1)) / 2;
    powerList(half_size+1:end) = [];

    powerListAll(i, :) = powerList(1:half_size);
end

load('metabolic_data.mat');

offset = 0.072; % (W/kg)

NoExo = metListAll(:, 1);
Assisted = metListAll(:, 4) - offset;

metDiff = NoExo - Assisted;

customColors = [
    0.1216, 0.4667, 0.7059;   % blue
    1.0000, 0.4980, 0.0549;   % orange
    0.1725, 0.6275, 0.1725;   % green
    0.8392, 0.1529, 0.1569;   % red
    0.5804, 0.4039, 0.7412;   % purple
    0.5490, 0.3373, 0.2941;   % brown
    0.8902, 0.4667, 0.7608;   % pink
    0.4980, 0.4980, 0.4980;   % gray
    0.7373, 0.7412, 0.1333;   % olive
    0.0902, 0.7451, 0.8118    % cyan
    ];

figure(); hold on;
xlabel('Exoskeleton Average Positive Mechanical Power (W/kg)');
ylabel('Metabolic Cost Reduction from No Exo (W/kg)')
for i = 1:10
    scatter(powerListAll(i, 3), metDiff(i), 80, ...
        'MarkerFaceColor', customColors(i, :), ...
        'MarkerEdgeColor', customColors(i, :), ...
        'MarkerFaceAlpha', 0.3, ...
        'MarkerEdgeAlpha', 0.9);
end

plot(mean(powerListAll(:, 3)), mean(metDiff(:)), 's', ...
    'MarkerEdgeColor', 'k', ...
    'MarkerFaceColor', 'k', ...
    'MarkerSize', 10, ...
    'HandleVisibility', 'off');

plot(mean(powerListAll(:, 3)), 0.72, 's', ...
    'MarkerEdgeColor', 'r', ...
    'MarkerFaceColor', 'r', ...
    'MarkerSize', 10, ...
    'HandleVisibility', 'off');

fileName = 'power_avg_metabolics_plot.svg';

if ~exist(fileName, 'file')
    saveas(gcf, fileName);
    fprintf('Saved: %s\n', fileName);
end
```
