## Supplementary material for "Experiment-free learning of exoskeleton assistance remains an unsolved problem": Replication Study Data and Code: plot_power_all.m

```
function plot_power_all(powerListAll)

groupNames = {'0% GC Shift', '5% GC Shift', '10% GC Shift', '15% GC Shift'};

subjectNumber = size(powerListAll, 1);
dataLength = size(powerListAll, 2);

meanVals = mean(powerListAll, 1);

figure(); hold on;

plot(NaN, NaN, 's', ...
    'MarkerEdgeColor', 'k', ...
    'MarkerFaceColor', 'k', ...
    'MarkerSize', 8, ...
    'DisplayName', 'Mean');

plot(NaN, NaN, 'o', ...
    'MarkerEdgeColor', 'k', ...
    'MarkerFaceColor', 'k', ...
    'MarkerSize', 6, ...
    'DisplayName', 'Individual Data');

% Boxplot
h = boxplot(powerListAll, ...
    'Colors', 'k', ...
    'Widths', 0.5, ...
    'Symbol', 'o', ...
    'MedianStyle', 'line', ...
    'Whisker', Inf);

lines = findall(gca, 'Type', 'Line');

for i = 1:length(lines)
    ls = get(lines(i), 'LineStyle');
    if strcmp(ls, '--')
        set(lines(i), 'LineStyle', '-', 'LineWidth', 0.5);
    end
end

boxColors = [ ...
    0.8500    0.3250    0.0980  % Red-orange (default MATLAB red-orange)
    0.9290    0.6940    0.1250  % Yellow (default MATLAB yellow)
    0.4940    0.1840    0.5560  % Purple (default MATLAB purple)
    0.4660    0.6740    0.1880  % Green (default MATLAB green)
    ];

boxes = findobj(gca, 'Tag', 'Box');
for k = 1:length(boxes)
    patch(get(boxes(k), 'XData'), get(boxes(k), 'YData'), ...
        boxColors(length(boxes)-k+1,:), ...
        'FaceAlpha', 0.3, 'EdgeColor', 'none', ...
        'HandleVisibility', 'off');
end

% Overlay mean and text
for k = 1:dataLength
    plot(k, meanVals(k), 's', ...
        'MarkerEdgeColor', 'k', ...
        'MarkerFaceColor', 'k', ...
        'MarkerSize', 8, ...
        'HandleVisibility', 'off');
end

customColors = [
    0.1216, 0.4667, 0.7059;   % blue
    1.0000, 0.4980, 0.0549;   % orange
    0.1725, 0.6275, 0.1725;   % green
    0.8392, 0.1529, 0.1569;   % red
    0.5804, 0.4039, 0.7412;   % purple
    0.5490, 0.3373, 0.2941;   % brown
    0.8902, 0.4667, 0.7608;   % pink
    0.4980, 0.4980, 0.4980;   % gray
    0.7373, 0.7412, 0.1333;   % olive
    0.0902, 0.7451, 0.8118    % cyan
    ];

spread = 0.2;
x_offsets = linspace(-spread, spread, subjectNumber);

for i = 1:subjectNumber
    scatter((1:dataLength) + x_offsets(i), powerListAll(i, :), 60, ...
        'MarkerFaceColor', customColors(i, :), ...
        'MarkerEdgeColor', customColors(i, :), ...
        'MarkerFaceAlpha', 0.7, ...
        'MarkerEdgeAlpha', 0.9, ...
        'HandleVisibility', 'off');
end

% Labels
ylabel('Mechanical Power (W kg^{-1})');
xlim([0.5, dataLength + 0.5]);
ylim([0, 0.3]);
xticks(1:dataLength);
xticklabels(groupNames);
legend('Location','best');

fileName = 'power_plot.svg';

if ~exist(fileName, 'file')
    saveas(gcf, fileName);
    fprintf('Saved: %s\n', fileName);
end

end
```
