## Supplementary material for "Experiment-free learning of exoskeleton assistance remains an unsolved problem": Replication Study Data and Code: plot_power_individual.m

```
function [] = plot_power_individual(fileNameList)

fileName = fileNameList{1};

parts = split(fileName);

pathName = parts{1};
subject = parts(1);

condName = cell(length(fileNameList), 1);
powerList = zeros(length(fileNameList), 1);

f1 = figure(); hold on; legend('Location', 'best');
xlabel('Gait Cycle (%)'); ylabel('Torque (Nm/kg)');

f2 = figure(); hold on; legend('Location', 'best');
xlabel('Gait Cycle (%)'); ylabel('Power (W/kg)');

defaultColor = [ ...
    0.0000    0.4470    0.7410  % Blue (default MATLAB blue)
    0.8500    0.3250    0.0980  % Red-orange (default MATLAB red-orange)
    0.9290    0.6940    0.1250  % Yellow (default MATLAB yellow)
    0.4940    0.1840    0.5560  % Purple (default MATLAB purple)
    0.4660    0.6740    0.1880  % Green (default MATLAB green)
    0.3010    0.7450    0.9330  % Light Blue (default MATLAB light blue)
    0.6350    0.0780    0.1840  % Dark Red (default MATLAB dark red)
    0.0000    0.5000    0.0000  % Green
    ];

for i = 1:length(fileNameList)
    fileName = fileNameList{i};
    parts = split(fileName);

    condName{i} = parts{3};
    matFileName = sprintf('%s Power.mat', fileName);
    matFullPath = fullfile(pathName, matFileName);

    load(matFullPath);

    switch pathName
        case 'S1', weight = 79; % [kg]
        case 'S2', weight = 106; % [kg]
        case 'S3', weight = 68; % [kg]
        case 'S4', weight = 54; % [kg]
        case 'S5', weight = 50; % [kg]
        case 'S6', weight = 69; % [kg]
        case 'S7', weight = 63; % [kg]
        case 'S8', weight = 58; % [kg]
        case 'S9', weight = 77; % [kg]
        case 'S10', weight = 70; % [kg]
    end

    cycle_steps = 1000;
    cycle = linspace(0, 100, cycle_steps)';

    torque_mean = mean(data_cycles.torque / weight, 2);
    power_mean = mean(data_cycles.power / weight, 2);

    colorIdx = i;

    figure(f1); hold on;
    plot(cycle, torque_mean, 'LineWidth', 1.5, ...
        'Color', defaultColor(colorIdx,:), 'DisplayName', condName{i});

    figure(f2); hold on;
    plot(cycle, power_mean, 'LineWidth', 1.5, ...
        'Color', defaultColor(colorIdx,:), 'DisplayName', condName{i});

    area(cycle, max(power_mean, 0), ...
        'FaceColor', defaultColor(colorIdx,:) * 0.6 + 0.4, ...
        'EdgeColor', 'none', ...
        'FaceAlpha', 0.3, ...
        'HandleVisibility', 'off');

    svgFileName = 'power_graph_plot.svg';

    if ~exist(svgFileName, 'file')
        saveas(gcf, svgFileName);
        fprintf('Saved: %s\n', svgFileName);
    end

    power_pos_avg_norm = mean(power_mean(power_mean > 0));
    power_neg_avg_norm = mean(power_mean(power_mean < 0));
    power_avg_norm = power_pos_avg_norm + power_neg_avg_norm;
    power_peak_norm = max(power_mean);
    
    save(matFullPath, 'data_cycles', 'power_pos_avg_norm', 'power_avg_norm', 'power_peak_norm');

    powerList(i) = power_pos_avg_norm;


end

figure(); hold on;
title(subject);
ylabel('Mechanical Power (W kg^{-1})');

barHandles = bar(powerList, 'HandleVisibility', 'off');

barHandles.FaceColor = 'flat';
for k = 1:length(powerList)
    colorIdx = min(k, length(powerList) - k + 1);

    barHandles.CData(k, :) = defaultColor(colorIdx, :);
end

xticks(1:length(fileNameList));
xticklabels(condName);
xtickangle(30);
end
```
