## Supplementary material for "Experiment-free learning of exoskeleton assistance remains an unsolved problem": Replication Study Data and Code: main_all.m

```
clear;

% N = 10
subjectNumList = [1, 2, 3, 4, 5, 6, 7, 8, 9, 10];

% N = 8
% subjectNumList = [1, 2, 3, 5, 6, 7, 8, 10];

for i = 1:length(subjectNumList)

    subjectNum = subjectNumList(i);
    fileNameList = generateFileNameList(subjectNum);

    metList = process_metabolics(fileNameList);

    half_size = (length(fileNameList)-1)/2;
    fileNameList = {fileNameList{1:half_size+1}};
    metList(2:half_size+1) = (metList(2:half_size+1) + metList(end:-1:half_size+2)) / 2;
    metList(half_size+2:end) = [];

    metListAll(i, :) = metList(2:half_size+1) - metList(1);
end

save('metabolic_data', 'metListAll');

plot_metabolics_all(metListAll);
```
