## Supplementary material for "Experiment-free learning of exoskeleton assistance remains an unsolved problem": Replication Study Data and Code: main_individual.m

```
clear;

subjectNum = 3;

fileNameList = generateFileNameList(subjectNum);

metList = process_metabolics(fileNameList);

plot_metabolics(fileNameList, metList);

fullLength = length(metList) - 1;

fileNameListAveraged = {fileNameList{1:fullLength/2+1}};
metListAveraged(1) = metList(1);
metListAveraged(2:1+fullLength/2) = (metList(2:1+fullLength/2) + metList(end:-1:2+fullLength/2)) / 2;

plot_metabolics(fileNameListAveraged, metListAveraged);
```
