## Supplementary material for "Experiment-free learning of exoskeleton assistance remains an unsolved problem": Replication Study Data and Code: plot_metabolics_all.m

```
function plot_metabolics_all(metListAll)

groupNames = {'No Exo', '0% GC Shift', '5% GC Shift', '10% GC Shift', '15% GC Shift'};

subjectNumber = size(metListAll, 1);
dataLength = size(metListAll, 2);

meanVals = mean(metListAll, 1);

baseline = meanVals(1);
percent_changes = (meanVals - baseline) / baseline * 100;

offset = 0.072; % [W/kg]
meanValsOffset = meanVals - offset;
percent_changes_2 = (meanValsOffset - baseline) / baseline * 100;

figure(); hold on;

plot(NaN, NaN, 's', ...
    'MarkerEdgeColor', 'k', ...
    'MarkerFaceColor', 'k', ...
    'MarkerSize', 8, ...
    'DisplayName', 'Mean');

plot(NaN, NaN, 's', ...
    'MarkerEdgeColor', 'b', ...
    'MarkerFaceColor', 'b', ...
    'MarkerSize', 8, ...
    'DisplayName', 'Mean (Weight Compensated)');

plot(NaN, NaN, 'o', ...
    'MarkerEdgeColor', 'k', ...
    'MarkerFaceColor', 'k', ...
    'MarkerSize', 8, ...
    'DisplayName', 'Individual Data');

% Boxplot
h = boxplot(metListAll, ...
    'Colors', 'k', ...
    'Widths', 0.5, ...
    'Symbol', 'o', ...
    'MedianStyle', 'line', ...
    'Whisker', Inf);

lines = findall(gca, 'Type', 'Line');

for i = 1:length(lines)
    ls = get(lines(i), 'LineStyle');
    if strcmp(ls, '--')
        set(lines(i), 'LineStyle', '-', 'LineWidth', 0.5);
    end
end

boxColors = [ ...
    0.8500    0.3250    0.0980  % Red-orange (default MATLAB red-orange)
    0.9290    0.6940    0.1250  % Yellow (default MATLAB yellow)
    0.4940    0.1840    0.5560  % Purple (default MATLAB purple)
    0.4660    0.6740    0.1880  % Green (default MATLAB green)
    0.3010    0.7450    0.9330  % Light Blue (default MATLAB light blue)
    ];

boxes = findobj(gca, 'Tag', 'Box');
for k = 1:length(boxes)
    patch(get(boxes(k), 'XData'), get(boxes(k), 'YData'), ...
        boxColors(length(boxes)-k+1,:), ...
        'FaceAlpha', 0.3, 'EdgeColor', 'none', ...
        'HandleVisibility', 'off');
end

% Overlay mean and text
for k = 1:dataLength
    plot(k, meanVals(k), 's', ...
        'MarkerEdgeColor', 'k', ...
        'MarkerFaceColor', 'k', ...
        'MarkerSize', 8, ...
        'HandleVisibility', 'off');

    text(k + 0.15, meanVals(k) + 0.05, ...
        sprintf('%.0f%%', percent_changes(k)), ...
        'HorizontalAlignment', 'center', ...
        'Color', 'black', 'FontSize', 10);

    if (k > 1)
        plot(k, meanValsOffset(k), 's', ...
            'MarkerEdgeColor', 'b', ...
            'MarkerFaceColor', 'b', ...
            'MarkerSize', 8, ...
            'HandleVisibility', 'off');

        text(k + 0.15, meanValsOffset(k) - 0.05, ...
            sprintf('%.0f%%', percent_changes_2(k)), ...
            'HorizontalAlignment', 'center', ...
            'Color', 'blue', 'FontSize', 10);
    end
end

dotColors = [
    0.1216, 0.4667, 0.7059;   % blue
    1.0000, 0.4980, 0.0549;   % orange
    0.1725, 0.6275, 0.1725;   % green
    0.8392, 0.1529, 0.1569;   % red
    0.5804, 0.4039, 0.7412;   % purple
    0.5490, 0.3373, 0.2941;   % brown
    0.8902, 0.4667, 0.7608;   % pink
    0.4980, 0.4980, 0.4980;   % gray
    0.7373, 0.7412, 0.1333;   % olive
    0.0902, 0.7451, 0.8118    % cyan
    ];

spread = 0.2;
x_offsets = linspace(-spread, spread, subjectNumber);

if subjectNumber == 8
    removedIndices = [4, 9];
    fullIndices = 1:10;
    keptIndices = setdiff(fullIndices, removedIndices);
    title('N = 8');
    svgFileName = 'metabolics_plot_n8.svg';
else
    keptIndices = 1:10;
    title('N = 10');
    svgFileName = 'metabolics_plot_n10.svg';
end

for i = 1:subjectNumber
    colorIdx = keptIndices(i);
    scatter((1:dataLength) + x_offsets(i), metListAll(i, :), 60, ...
        'MarkerFaceColor', dotColors(colorIdx, :), ...
        'MarkerEdgeColor', dotColors(colorIdx, :), ...
        'MarkerFaceAlpha', 0.7, ...
        'MarkerEdgeAlpha', 0.9, ...
        'HandleVisibility', 'off');
end

% Labels
ylabel('Metabolic Cost (W/kg)');
xlim([0.5, dataLength + 0.5]);
ylim([2, 5.5]);
xticks(1:dataLength);
xticklabels(groupNames);
xtickangle(0);
legend('Location','best');

if ~exist(svgFileName, 'file')
    saveas(gcf, svgFileName);
    fprintf('Saved: %s\n', svgFileName);
end
```
