## Supplementary material for "Experiment-free learning of exoskeleton assistance remains an unsolved problem": Replication Study Data and Code: plot_metabolics_individual.m

```
function [] = plot_metabolics_individual(fileNameList, metList)

parts = split(fileNameList{1});

figure(); hold on;
title(parts(1));
ylabel('Metabolic Cost (W kg^{-1})');

defaultColor = [ ...
    0.0000    0.4470    0.7410  % Blue (default MATLAB blue)
    0.8500    0.3250    0.0980  % Red-orange (default MATLAB red-orange)
    0.9290    0.6940    0.1250  % Yellow (default MATLAB yellow)
    0.4940    0.1840    0.5560  % Purple (default MATLAB purple)
    0.4660    0.6740    0.1880  % Green (default MATLAB green)
    0.3010    0.7450    0.9330  % Light Blue (default MATLAB light blue)
    0.6350    0.0780    0.1840  % Dark Red (default MATLAB dark red)
    0.0000    0.5000    0.0000  % Green
    0.7500    0.0000    0.7500  % Purple
    0.7500    0.7500    0.0000  % Olive
    0.0000    0.7500    0.7500  % Teal
    ];

barHandles = bar(metList(2:end) - metList(1), 'HandleVisibility', 'off');

barHandles.FaceColor = 'flat';
for k = 1:length(metList)-1
    barHandles.CData(k, :) = defaultColor(k+1, :);
end

condName = cell(length(fileNameList)-1, 1);

for i = 2:length(fileNameList)
    parts = split(fileNameList{i});
    condName{i-1} = strjoin(parts(3:end), ' ');
end

xticks(1:length(fileNameList)-1);
xticklabels(condName);
xtickangle(30);

metBaseLine = metList(2);

percent_changes = (metList(2:end) - metBaseLine) / (metBaseLine - metList(1)) * 100;

yline(metBaseLine - metList(1), 'Color', defaultColor(1, :), 'HandleVisibility', 'off');

for i = 1:length(percent_changes)
    text(i, metList(i+1) - metList(1), sprintf('%.1f%%', percent_changes(i)), ...
        'VerticalAlignment', 'bottom', 'HorizontalAlignment', 'center', 'Color', 'red');
end

end
```
