## Supplementary material for "Experiment-free learning of exoskeleton assistance remains an unsolved problem": Replication Study Data and Code: process_metabolics.m

```
function metList = process_metabolics(fileNameList)

parts = split(fileNameList{1});

averageTime = 2; % [min]

metList = zeros(length(fileNameList), 1);

for i = 1:length(fileNameList)

    fileName = fileNameList{i};

    pathName = parts{1};

    switch pathName
        case 'S1', weight = 79; % [kg]
        case 'S2', weight = 106; % [kg]
        case 'S3', weight = 68; % [kg]
        case 'S4', weight = 54; % [kg]
        case 'S5', weight = 50; % [kg]
        case 'S6', weight = 69; % [kg]
        case 'S7', weight = 63; % [kg]
        case 'S8', weight = 58; % [kg]
        case 'S9', weight = 77; % [kg]
        case 'S10', weight = 70; % [kg]
    end

    T = readcell(strcat(fullfile(pathName,fileName),'.xlsx'));

    % Find the index of the columns containing EEm, VO2, and VCO2
    time_index = find(strcmp(T(1,:),'t'));
    VO2_index = find(strcmp(T(1,:),'VO2'));
    VCO2_index = find(strcmp(T(1,:),'VCO2'));
    
    time = cell2mat(T(4:end, time_index))*86400; % [s] 1 day = 86400 seconds

    time = time - time(1);

    VO2 = cell2mat(T(4:end, VO2_index)); % [mL/min]
    VCO2 = cell2mat(T(4:end, VCO2_index)); % [mL/min]

    y_meas = (0.278*VO2 + 0.075*VCO2)/weight; % [W/kg]

    startTime = time(end) - 60 * averageTime;
    [~, startTimeIdx] = min(abs(time - startTime));

    y_average = mean(y_meas(startTimeIdx:end));

    metList(i) = y_average;

    fprintf('%d: (%ds) %s = %.4f W/kg\n', i, round(time(end)), fileName, y_average);
end

end
```
