## Supplementary material for "Experiment-free learning of exoskeleton assistance remains an unsolved problem": Replication Study Data and Code: README.md

This repository contains MATLAB scripts and data for analyzing **mechanical power** and **metabolic cost** in participants using an exoskeleton device. The analysis includes both **individual-level** and **group-level** comparisons across different experimental conditions.

## Requirements

- MATLAB Version: **R2024a (24.1.0.2689473) Update 6**
- Required Toolbox:  
  - **Statistics and Machine Learning Toolbox**

---

### Experimental Conditions

Participants completed a total of **10 walking trials**, consisting of 5 experimental conditions repeated in a **double-reversal design** to minimize order effects. Each trial lasted **5 minutes** at a fixed treadmill speed of **1.25 m/s**.

The experimental conditions were:

- **C0**: No Exoskeleton (Baseline)
- **C1**: 0% Gait Cycle Shift
- **C2**: 5% Gait Cycle Shift
- **C3**: 10% Gait Cycle Shift
- **C4**: 15% Gait Cycle Shift

Conditions C1–C4 were presented in a **randomized order**, and the entire set of five conditions was repeated in reverse.

**Reference Torque**
- `Applied torque.xlsx`

**Participant's Information**
- `Demographic information.xlsx`  
  Contains participant information:
  - Gender
  - Age  
  - Height  
  - Weight  
  - Condition order  
  
---

## Data Structure

### **Exoskeleton (Exo) Data**

#### `Exo Data/`
- **Raw Data (.mat file):**
  - `data_cycles` (structure):
    - `Time (s)`  
    - `gaitCycle (%)`  
    - `torque (Nm)`: (-) Extension, (+) Flexion 
    - `power (W)`  
  - Summary Metrics:
    - `power_avg_norm (W/kg)` – average normalized power  
    - `power_pos_avg_norm (W/kg)` – average of positive normalized power  
    - `power_peak_norm (W/kg)` – peak normalized power  

#### Scripts

- `main_individual.m` (`.mlx` file includes figure in the Live Script output)  
  → Visualize **torque** and **power** over the gait cycle **within each participant**  
  → Calls: `plot_power_individual.m`  
  → Output: `power_graph_plot.svg`

- `main_all.m` (`.mlx` file includes figure in the Live Script output)  
  → Compare **positive mechanical power** of all pariticipants **across conditions**  
  → Calls: `plot_power_all.m`  
  → Output: `power_plot.svg`

- `main_average_power_metabolics.m` (`.mlx` file includes figure in the Live Script output)  
  → Compare **average power** and **metabolic cost**  
  → Output: `power_avg_metabolics_plot.svg`

- `generateFileNameList.m`  
  → Helper function to generate standardized filenames

---

### **Metabolic Data**

#### `Metabolic Data/`

- **Raw Data**:  
  - `.xlsx` file from **COSMED K5** metabolic system  

#### Scripts

- `main_individual.m` (`.mlx` file includes figure in the Live Script output)  
  → Compare **metabolic cost across conditions** **within each participant**  
  → Calls: `plot_metabolics_individual.m`  
  → Includes **double-reversal** result analysis

- `main_all.m` (`.mlx` file includes figure in the Live Script output)  
  → Compare **metabolic cost across all participants** and conditions  
  → Calls: `plot_metabolics_all.m`  
  → Output:  
    - `metabolics_plot_n10.svg` (10 participants)  
    - `metabolics_plot_n8.svg` (8 participants)

- `process_metabolics.m`  
  → Process raw metabolic `.xlsx` files

- `generateFileNameList.m`  
  → Helper function to generate standardized filenames
