## Supplementary figures and images for "Experiment-free learning of exoskeleton assistance remains an unsolved problem"

### metabolics_plot_n8.png

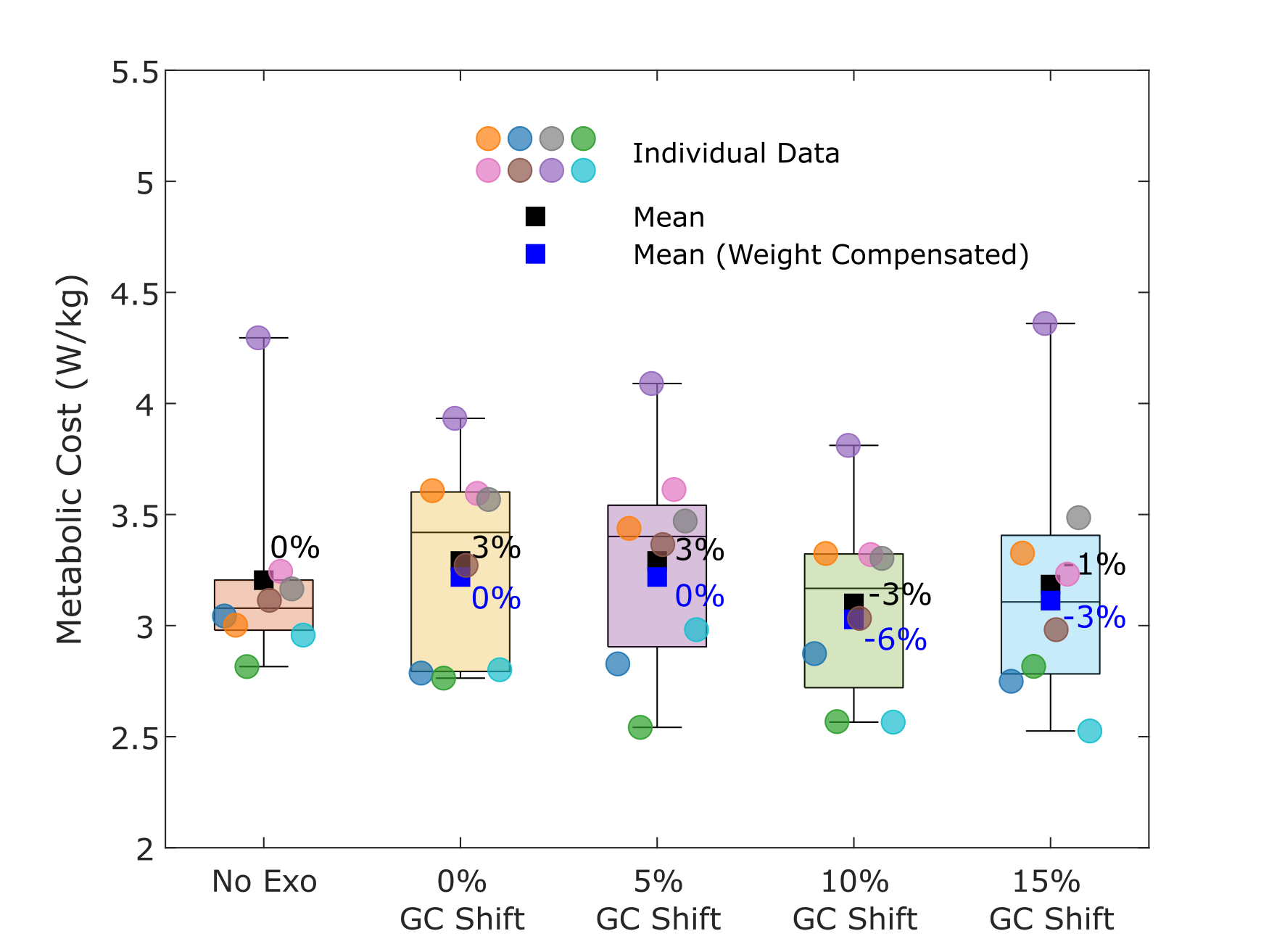

### metabolics_plot_n10.png

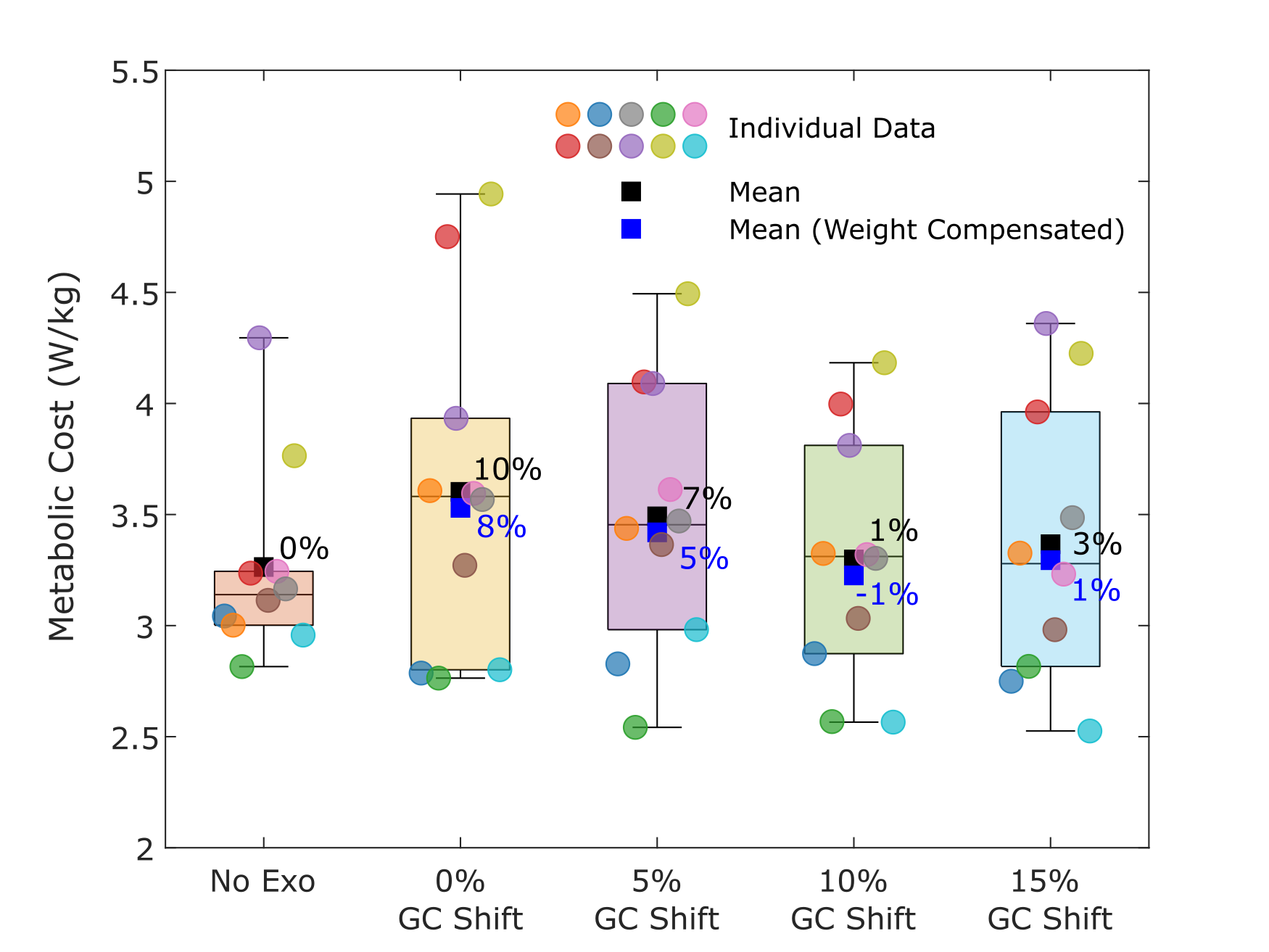

### power_avg_metabolics_plot.png

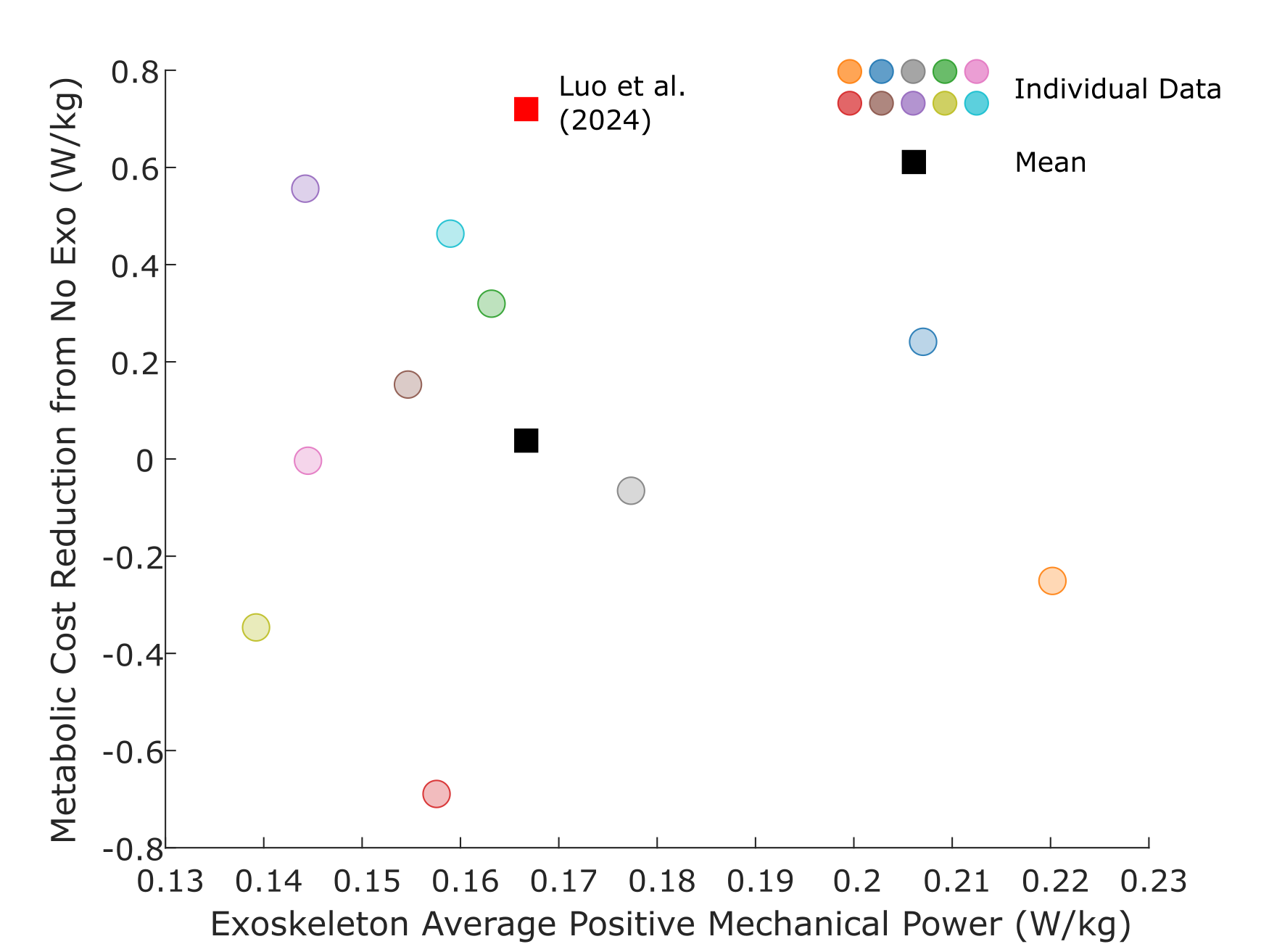

### power_graph_plot.png

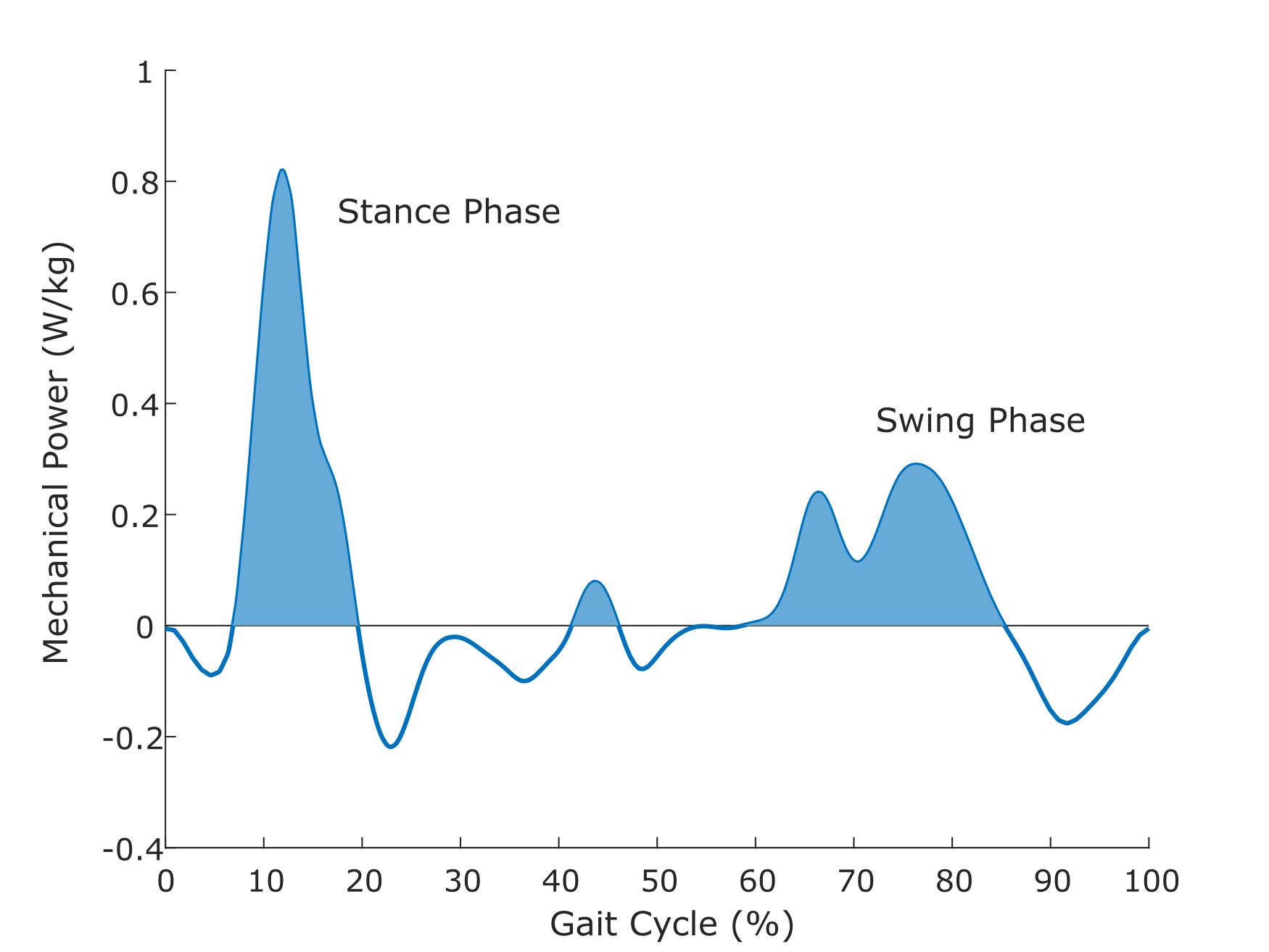

### power_plot.png

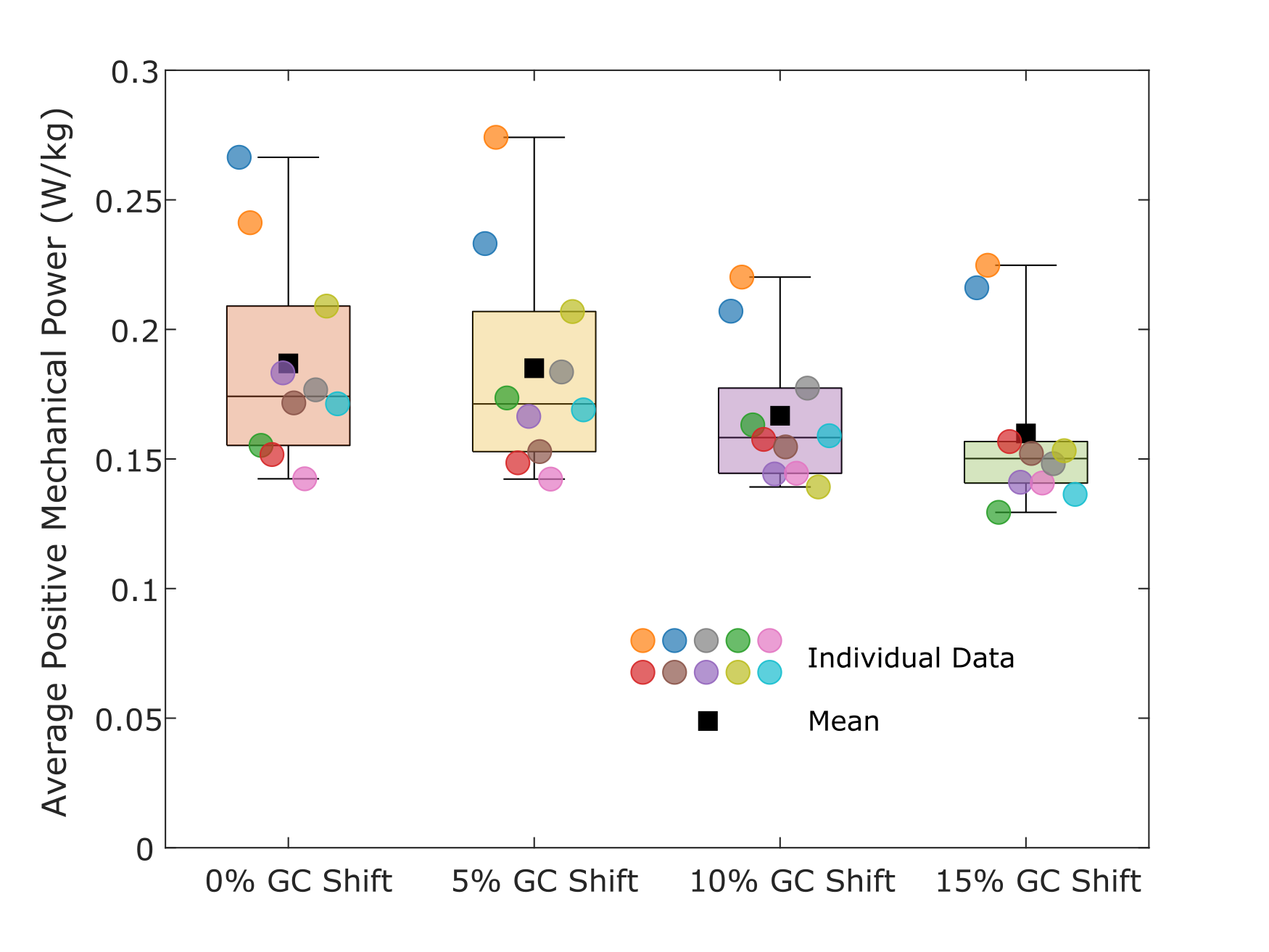
